# Photoperiodic Plasticity of Transcriptomic and Photosynthetic Rhythms in the Liverwort *Marchantia polymorpha*

**DOI:** 10.64898/2026.10.20.760308

**Authors:** Marcos Ramos-González, M. Elena García-Gómez, Jorge Domíguez-Barragán, Mercedes García-González, Francisco J. Romero-Campero

## Abstract

Earth’s translation around the Sun and tilted rotational axis drive periodic seasonal changes in day length, or photoperiod, in temperate zones. In photosynthetic organisms, light acts not only as an environmental cue but also as the primary energy source. Therefore, characterizing the plasticity of oscillating molecular systems during photoperiod acclimation is essential for understanding the temporal orchestration of physiological processes in these organisms. Although photoperiodic plasticity has been extensively studied in angiosperms and microalgae, its impact on the molecular rhythms in non-vascular plants (bryophytes) remains poorly understood. In this study, we performed an integrative temporal analysis combining transcriptomic and photosynthetic approaches in the non-vascular model plant *Marchantia polymorpha*. Plants were grown under long (summer) and short (winter) photoperiods and subsequently transferred to constant light or darkness to unravel the light/dark-responsive and circadian components of their rhythmic transcriptome profiles. Plants under short photoperiod exhibited a higher number of rhythmic genes and an advance in their expression pattern, affecting several pathways including photosynthesis. However, only ∼25% of these genes remained oscillating under constant conditions. We further identified dusk responsiveness and dawn anticipation as key features of the rhythmic patterns in *Marchantia polymorpha*. These patterns were associated with specific transcription factors, highlighting a potential role for *MpHY5* in circadian regulation in this liverwort. Consistent with the transcriptomic results, analyses of photosynthetic physiology and chloroplast photorelocation revealed a general anticipation under short photoperiods and mechanisms mitigating photooxidative stress at dawn in both photoperiods.

## Introduction

The occurrence of molecular rhythmic patterns underpinning physiological processes is widespread across living organisms. These physiological rhythms are associated with adaptation to pervasive cyclical environmental cues. Among these, Earth’s rotation plays a key role by generating 24-hour light/dark cycles, known as diel cycles. In turn, Earth’s tilted rotational axis and translation around the Sun produce consistent seasonal changes in environmental cues such as temperature and day length or photoperiod in temperate regions. Given that light constitutes the primary energy source for photosynthetic organisms, it acts as a decisive environmental signal, shaping seasonal acclimation to diel rhythmicity, from long summer days (LD, 16h light : 8h dark) to short winter days (SD, 8h light : 16h dark). Since these environmental variations are predictable, organisms do not only respond to them, but have developed molecular systems that anticipate them to maximize organismal fitness. These systems are referred to as circadian clocks. At the transcriptional level, the core of circadian clocks is constituted by networks of transcription factors that control their own expression through feedback loops. These gene networks generate autonomous self-sustained oscillations that are entrained by external cues such as light (Nakamichi 2011). Circadian clocks orchestrate the acclimation to different photoperiods by temporarily segregating the occurrence at specific time points of physiological processes like photosynthesis, protein synthesis, carbon fixation, and DNA replication (Zones et al. 2015; Romero-Losada et al. 2025). Some core components in circadian clocks are conserved across the green lineage or *Viridiplantae*. Nevertheless, the number of gene copies varies considerably between species, leading to substantial differences in network topology (Laosuntisuk et al. 2023). Despite these differences in circadian clock architecture, diel transcriptional programs orchestrating biological processes display similarities across diverse species in the green lineage (Ferrari et al. 2019; Tan et al. 2024). Diel transcriptional programs combine direct responses to light and darkness with autonomous environment-independent circadian regulation. These two components can be discriminated by analysing rhythmicity under free-running conditions consisting of constant light or darkness (Romero-Losada et al. 2025).

Bryophytes alongside tracheophytes constitute the entirety of extant land plants. Although bryophytes predate tracheophytes evolutionary, non-vascular plants possess a larger gene family space than their sister clade of vascular plants. This has contributed to their adaptation to a wide range of extreme environments (Dong et al. 2025). Within non-vascular plants, the liverwort *Marchantia polymorpha* (Marchantia) stands out due to its relative reduced genome size compared to other land plants, resulting from the absence of ancient whole-genome duplications (Bowman et al. 2022). Consequently, Marchantia has become a widely used model in evolutionary studies, particularly those aimed at elucidating the molecular mechanisms that promoted land colonization by plants (Catarino et al. 2026). In chronobiology, Marchantia is especially interesting because it lacks a *CIRCADIAN CLOCK ASSOCIATED 1 / LATE ELONGATED HYPOCOTYL* (*CCA1/LHY*) orthologue, a ubiquitous component of circadian clocks in the green lineage from marine microalgae to terrestrial plants (Linde et al. 2017). However, homologous proteins to those found in the *Arabidopsis thaliana* (Arabidopsis) clock, such as *PSEUDO-RESPONSE REGULATOR* (*PRR*), *TIMING OF CAB EXPRESSION 1* (*TOC1*), *REVEILLE* (*RVE*), *EARLY FLOWERING 3* (*ELF3*), *GIGANTEA* (*GI*), *LUX ARRHYTHMO* (*LUX*) and *DE-ETIOLATED 1* (*DET1*), have been identified in this liverwort. Among these, a conserved transcriptional feedback loop constituted by *MpPRR*, *MpRVE* and *MpTOC1* has been suggested (Linde et al. 2017; Lagercrantz et al. 2020). These proteins exhibit rhythmic expression under free-running conditions, making them good candidates to generate self-sustained circadian expression oscillations (Linde et al. 2017; Lagercrantz et al. 2021; Petersen et al. 2022). Most of these regulators have been studied based on their homology with the Arabidopsis circadian clock, where its components are organized into groups according to their expression phase or peaking time point. Thus, in Arabidopsis, the Evening Complex (EC) constituted by *LUX*, *ELF3* and *ELF4*, present orthologs in Marchantia. The expression of PRRs in Arabidopsis extends throughout the day, with *PRR9* peaking in the morning, *PRR5* and *PRR7* around midday, and *TOC1* and *PRR3* toward the end of the light period (Farré and Liu 2013). In Marchantia, *MpPRR* phylogenetically groups within the Arabidopsis *PRR7/3* clade and *MpTOC1* clusters with *TOC1/PRR1* (Hotta 2022). No early morning component has yet been identified in the Marchantia clock. Nevertheless, the existence of autonomous *MpDET1* rhythms, which in Arabidopsis are dependent on *CCA1* rhythms, suggests a relevant role for this protein in the Marchantia circadian clock that still requires further characterization (Lagercrantz et al. 2021). The reduced size of the Marchantia circadian clock may account for the weaker and more rapidly dampened rhythms observed under free-running conditions compared to angiosperms (Linde et al. 2017). Light receptors also differ substantially in copy number from those characterized in flowering plants. Marchantia possesses single copy gene encoding the photoreceptors phytochrome (*MpPHY*), cryptochrome (*MpCRY*), UV RESISTANCE LOCUS8 (*MpUVR8*), and phototropin (*MpPHOT*), as well as key signaling proteins such as *PHYTOCHROME INTERACTING FACTOR* (*MpPIF*) (Komatsu et al. 2014; Inoue et al. 2016, 2019). Due to the lack of redundancy, these proteins differ from their counterparts in Arabidopsis where they are part of multigene families. For example, *MpPHY* displays characteristics from type I and type II phytochromes, responding to both red and far-red irradiance (Inoue et al. 2016). Similarly, *MpPHOT* exhibits photoactivation properties resembling those of phot2-type receptor together with thermochemical proporties characteristic of phot1-type receptors (Komatsu et al. 2014; Kato et al. 2021).

Despite this progress, studies conducted to date have largely overlooked the effect of photoperiod. This environmental cue has been described as one of the main entrainment signals of the circadian clock in multiple plant species (Flis et al. 2016; Xiang et al. 2022; Romero-Losada et al. 2025). In Marchantia, photoperiod regulates key reproductive and developmental processes. Specifically, LD conditions promote the formation of antheridiophores and archegoniophores as well as incresing total plant size, whereas SD conditions increase the number of gemmae cups (Voth and Hamner 1940; Yamaoka et al. 2021; Biswal and Panigrahi 2025). In addition, chloroplast-encoded genes involved in light-independent chlorophyll biosynthesis have been shown to play an important role under SD conditions, suggesting that photoperiod may have influenced the evolutionary retention of these genes (Ueda et al. 2014). In contrast to Arabidopsis, photoperiod length measurement in Marchantia appears to depend primarily on the relative light/dark proportion in diel cycles, with a more limited contribution of the circadian clock (Kanesaka et al. 2023). Furthermore, in Arabidopsis, twilight length has recently been identified as a key determinant for sensing the seasonal context throught mechanisms mediated by *CCA1/LHY* (Mehta et al. 2026). The absence of these proteins in Marchantia, together with its distinct photoperiodic regulation and weaker circadian rhythmicity, raises important questions regarding the impact of photoperiod on its diel transcriptional programs.

Light harvesting is one of the major physiological processes regulated by diel cycles. A key determinant of light harvesting is the chloroplast area exposed at the cell surface. Therefore, most plants, including Marchantia, reorganize the intracellular position of their chloroplasts depending on the light conditions through a process known as photorelocation. This process involves movement between periclinal (cell surface) and anticlinal (sidewall) cell walls (Komatsu et al. 2014). Under weak blue light, chloroplasts accumulate at the periclinal wall, whereas strong blue light induces avoidance movement toward anticlinal walls. In darkness, chloroplasts also display characteristic positioning in anticlinal walls (Komatsu et al. 2014). Photorelocation depends on short actin filaments surrounding chloroplasts (Suetsugu and Wada 2016). This cellular process does not appear to be ancestral in the green lineage or Viridiplantae, having emerged after the divergence between Chlorophyta and Streptophyta (Suetsugu and Wada 2016). This likely reflects the ability of microalgae to regulate light exposure through vertical movement in the water column, whereas terrestrial plants are sessile organisms that must acclimate to fluctuating light conditions while attached to a substrate (Wada 2005).

Therefore, despite the growing interest in Marchantia oscillating molecular systems, the relative contributions of light/dark-responsiveness and environment-independent circadian regulation under different photoperiods remain largely unexplored. This study characterizes the acclimation to LD and SD conditions of transcriptomic and photosynthetic rhythms in Marchantia. Environment-independent circadian rhythms are disentangle from light/dark-responsiveness by including free-running conditions consisting of constant light (LL) and constant darkness (DD) following entrainment under both photoperiods. Our results revealed an orchestration between light/dark responsive processes and circadian anticipation in shaping rhythmic transcriptome dynamics underpinning the coordination of photosynthetic acclimation to seasonal changes in daylength in this liverwort.

## Results

### Rhythmic gene expression under diel cycles and constant conditions

Marchantia plants were cultivated under light/dark cycles at a constant temperature under LD (16 h light:8 h darkness) and SD conditions (8 h light:16 h darkness) (Sup. Fig. S1). Samples were collected every 4 h from the onset of the light period, denoted by Zeitgeber Time 0 (ZT0). Principal Component Analysis revealed a temporal clustering of the global transcriptomic profiles forming cyclic closed trajectories capturing diel cycles (Sup. Fig. S2).

Photoperiod can affect the number of rhythmic transcripts in diel cycles (de los Reyes et al. 2017; Wu et al. 2023). In Marchantia, photoperiod presented a marked effect on transcriptome rhythmicity, with 7,048 rhythmic genes identified under LD compared to 9,130 under SD conditions. Among them, 5,937 genes exhibited rhythmicity regardless of the photoperiod of entrainment (**Fig. 1A**). This observation is consistent with previous studies assessing photoperiodic effects on gene expression rhythms in plants (do Nascimento et al. 2025; Romero-Losada et al. 2025). A significant 100-fold increase in the expression levels of rhythmic genes was detected compared to non-rhythmic genes (**Fig. 1B**). Analysis of the expression levels of the 18,201 annotated genes revealed that 5,523 did not reach a gene expression threshold of 2 FPKM (Fragments Per Kilobase of transcript and Million mapped reads) in any sample. This group showed significant enrichment in microtubule-based movement, DNA damage checkpoint signaling, response to oxidative stress, protein phosphorylation, negative regulation of cell cycle, epigenetic regulation of gene expression and gene silencing by RNA (**Fig. 1C**). Therefore, genes with little or no expression are significantly associated with specific stress responses and could potentially exhibit rhythmicity under the specific stress conditions inducing in them higher expression levels. Similar observations have been reported for multiple stresses, whose associated transcripts exhibit rhythmic expression through the diel cycle in agreement with the expected phase of the corresponding stress (Avelino et al. 2025). Hereinafter, all calculations will be performed using the number of genes expressed in at least one of the samples (12,678 genes).

**Figure 1.**
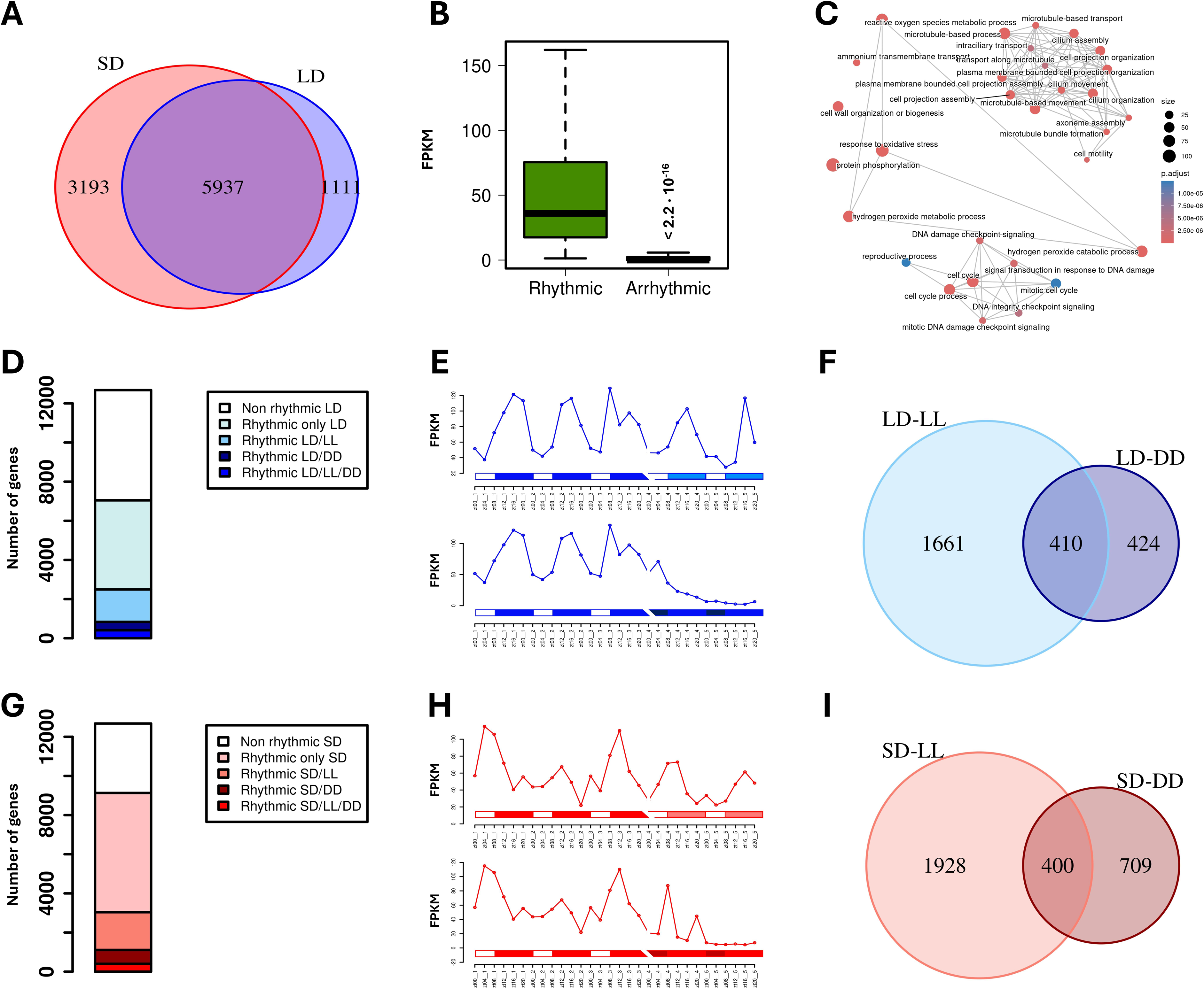
Transcriptome rhythmicity under alternating light/dark cycles and free-running conditions. **A)** Venn diagram comparing rhythmic genes under LD (blue) and SD (red). **B)** Boxplot representing expression of rhythmic (green) and arrhythmic (blue) genes. Gene expression levels are measured as FPKM. Significance was computed according to Mann–Whitney–Wilcoxon nonparametric test. **C)** Association plot showing significant GO BP terms enrichment for low-expressed genes (< 2 FPKM) in all samples. **D)** Stacked bar plot representing the total number of rhythmic genes under different conditions and LD-entrainment. **E)** Example of the expression profile of an LD-entrained-LL-rhythmic gene whose expression dampens progressively under DD. White rectangles represent light periods, blue-filled rectangles correspond dark periods under LD. ZTN, Zeitgeber time N, marks the time point N hours after dawn (lights on, ZT0). CTN, circadian time N denotes the time point N hours after subjective dawn. Light blue-filled rectangles represent subjective night under LL and dark blue-filled rectangles represent subjective day under DD. A discontinuity is shown on the time axis to indicate that samples were collected after 24 h acclimation to the corresponding free-running conditions. **F)** Venn diagram comparing LL-(light blue) and DD-rhythmic (dark blue) genes after LD entrainment. **G)** Stacked bar plot representing the total number of rhythmic genes under different conditions and SD-entrainment. **H)** Example of the expression profile of a SD-entrained-LL-rhythmic gene whose expression dampens progressively under DD. White rectangles represent light periods, red-filled rectangles correspond dark periods under SD. Light red-filled rectangles represent subjective night under LL and dark red-filled rectangles represent subjective day under DD. A discontinuity is shown on the time axis to indicate that samples were collected after 24 h acclimation to the corresponding free-running conditions. **I)** Venn diagram comparing LL-(light red) and DD-rhythmic (dark red) genes after SD entrainment.

Gene expression rhythms may result from circadian control or responses to light/dark cycles (Mehta et al. 2021). To distinguish between circadian and light/dark-responsive rhythmic genes, plants were transferred to free-running conditions, consisting of either constant light (LL) or constant darkness (DD). Plants were acclimated for one day to free-running conditions and sampling began at subjective dawn (CT0, Circadian Time 0) on the following day (Sup. Fig. S1). A clear dependence on the entraining photoperiod was detected in the proportion of genes maintaining rhythmicity under free-running conditions, despite a marked overall reduction in transcriptome rhythmicity under these conditions. In LD-entrained plants, 2,071 genes remained rhythmic after transfer to LL conditions, whereas 834 maintained rhythmicity under DD conditions (**Fig. 1D**). Following SD entrainment, these numbers increased to 2,328 and 1,109, respectively (**Fig. 1G**). Among these, 410 genes remained rhythmic under both LL and DD conditions following LD entrainment (**Fig. 1F**), whereas 400 rhymtic genes were identified following SD entrainment (**Fig. 1I**). Therefore, the detrimental effect of DD on transcriptome rhythmicity was substantially stronger than that of LL, regardless of the photoperiodic entrainment regime (**Fig. 1E,H**), similarly to previous observations in plants like *A. thaliana* (Anwer et al. 2014) and opposite to results reported in marine phytoplankton like *Ostreococcus tauri* (Romero-Losada et al. 2025).

Only a small core of 45 genes remained rhythmic under all conditions (**Fig. 2A**), hereafter referred to as bona fide circadian genes. Interestingly, the number of genes maintaining rhythmicity under both LL and DD conditions was higher in LD-entrained plants, despite the greater number of free-running rhythmic genes detected after SD entrainment. Notably, the set of free-running rhythmic genes was strongly influenced by the photoperiod of entrainment, with only 10% shared between both photoperiods (**Fig. 2A**). To assess the association of these gene groups with specific biological processes, functional enrichment analysis was performed for LL- and DD-rhythmic genes under both photoperiods. No functional enrichment was detected for DD-rhythmic genes, whereas LL-rhythmic genes were significantly associated with processes related to photosynthesis, the biosynthesis of organic acids and isoprenoids, including carotenoids (**Fig. 2B**). The photoreceptor *MpPHOT* (Mp5g03810) and the transcription factor *MpPIF* (Mp3g17350), involved in the control of processes associated with photosynthesis and light capture (Komatsu et al. 2014; Inoue et al. 2016, 2019), were identified among the LL-rhythmic but DD-arrhythmic genes. Both genes presented photoperiodic variations in their rhythmic profiles. In contrast, the photoreceptors *MpCRY* (Mp2g17590) and *MpUVR8* (Mp2g11590) required alternating light/dark cylces to maintain rhythmicity, whereas the photo-thermo receptor *MpPHY* (Mp2g16090) was rhythmic exclusively under SD conditions. Bona fide circadian genes were enriched in processes related to chloroplast protein localization and membrane lipid biosynthesis such as ceramides (**Fig. 2C**). Regarding transfers to LL after entrainment under different photoperiods, LD-LL-specific-rhythmic genes were enriched in tRNA aminoacylation for protein translation, steroid biosynthesis, plastid organization and phosphate ion transport (Sup. Table S1). In contrast, SD-LL-specific-rhythmic genes showed enrichment in photosynthesis and tetrapyrrole biosynthesis such as chlorophyll (Sup. Table S2). These results suggest a tight circadian regulation of photosynthetic and chloroplast associated processes together with a seasonal acclimation of this regulation exerted by the photoperiod.

**Figure 2.**
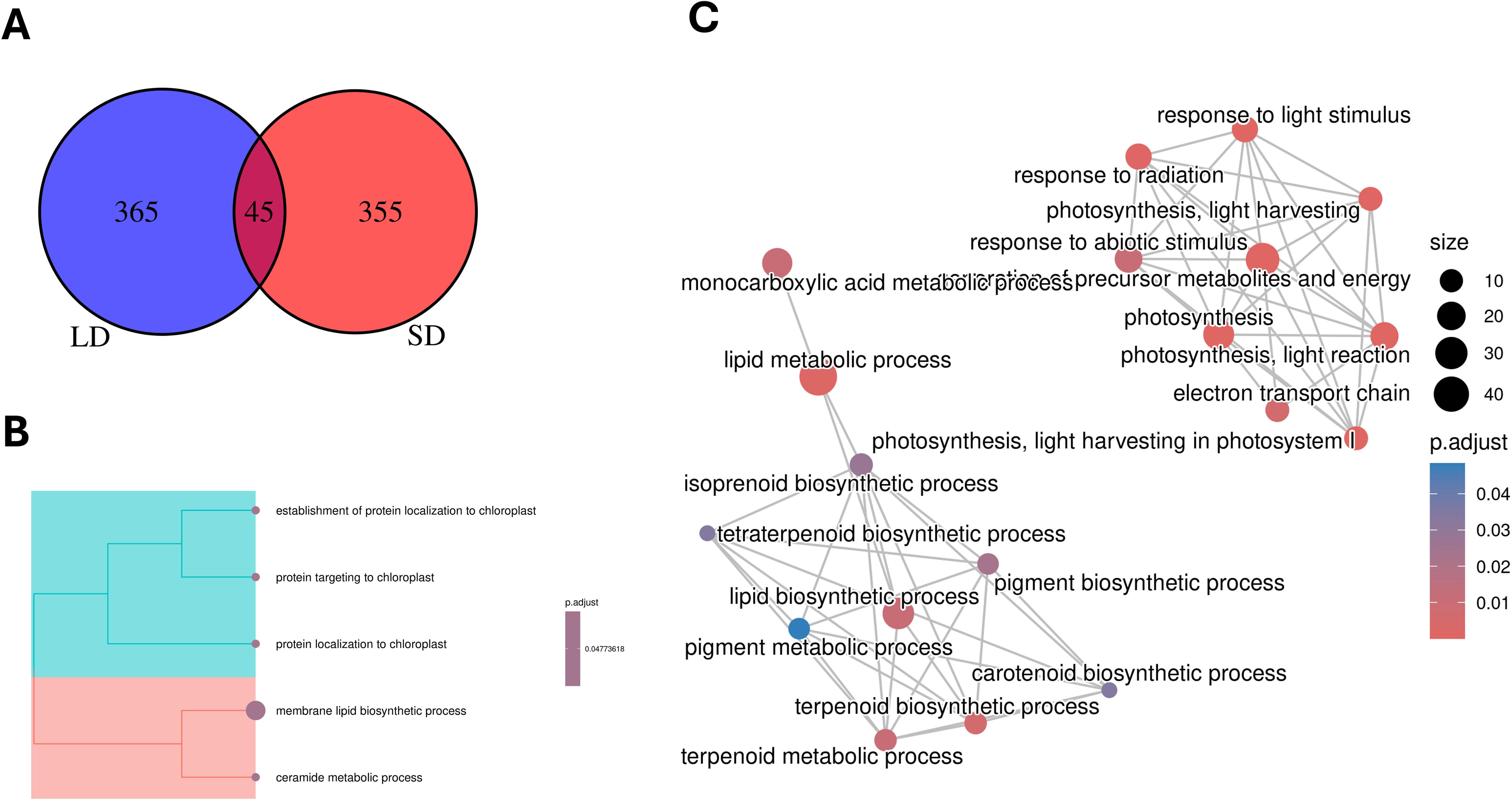
Identification of bona fide-circadian genes under each photoperiod. **A)** Venn diagram depicting the total number of circadian genes after LD (blue) or SD entrainment (red). **B)** Treeplot showing significant GO BP terms enrichment for rhythmic genes under all tested conditions. **C)** Association plot showing significant GO BP terms enrichment for LL-rhythmic genes under both photoperiods.

In addition to the overall loss of rhythmicity identified under free-running conditions, specific changes in the wave parameters of genes maintining rhythmicity were detected in agreement with previous studies (Ueno et al. 2022; Romero-Losada et al. 2025). In Marchantia, transfer to free-running conditions caused a decrease in the amplitude of LL- and DD-rhythmic genes for both photoperiods of entrainment (**Fig. 3A**), an effect previously attributed to desynchronization between the autonomous circadian clocks of individual cells (Wenden et al. 2012). Between 74 and 87% of rhythmic genes exhibited reduced amplitude, with a slightly stronger effect observed in LD-entrained plants suggesting a higher synchronization in SD-entrained plants that were less affected when transferred to free-running conditions. The wave mesor, corresponding to the mean of the oscillatory expression level, was only significantly affected under DD conditions after SD entrainment, resulting in rhythmic genes displaying higher mesor values than under light/dark cycles (**Fig. 3B**) indicating a certain level of activation by dark in these genes that could support their oscillations under DD conditions. Comparison between LL- and DD-rhythmic gene sets further revealed significantly lower mesor values under DD than under LL independently from the photoperiod of entrainment. However, this reduction in expression was not observed in the sets of rhythmic non-circadian genes under either photoperiod (Sup. Fig. S3).

**Figure 3.**
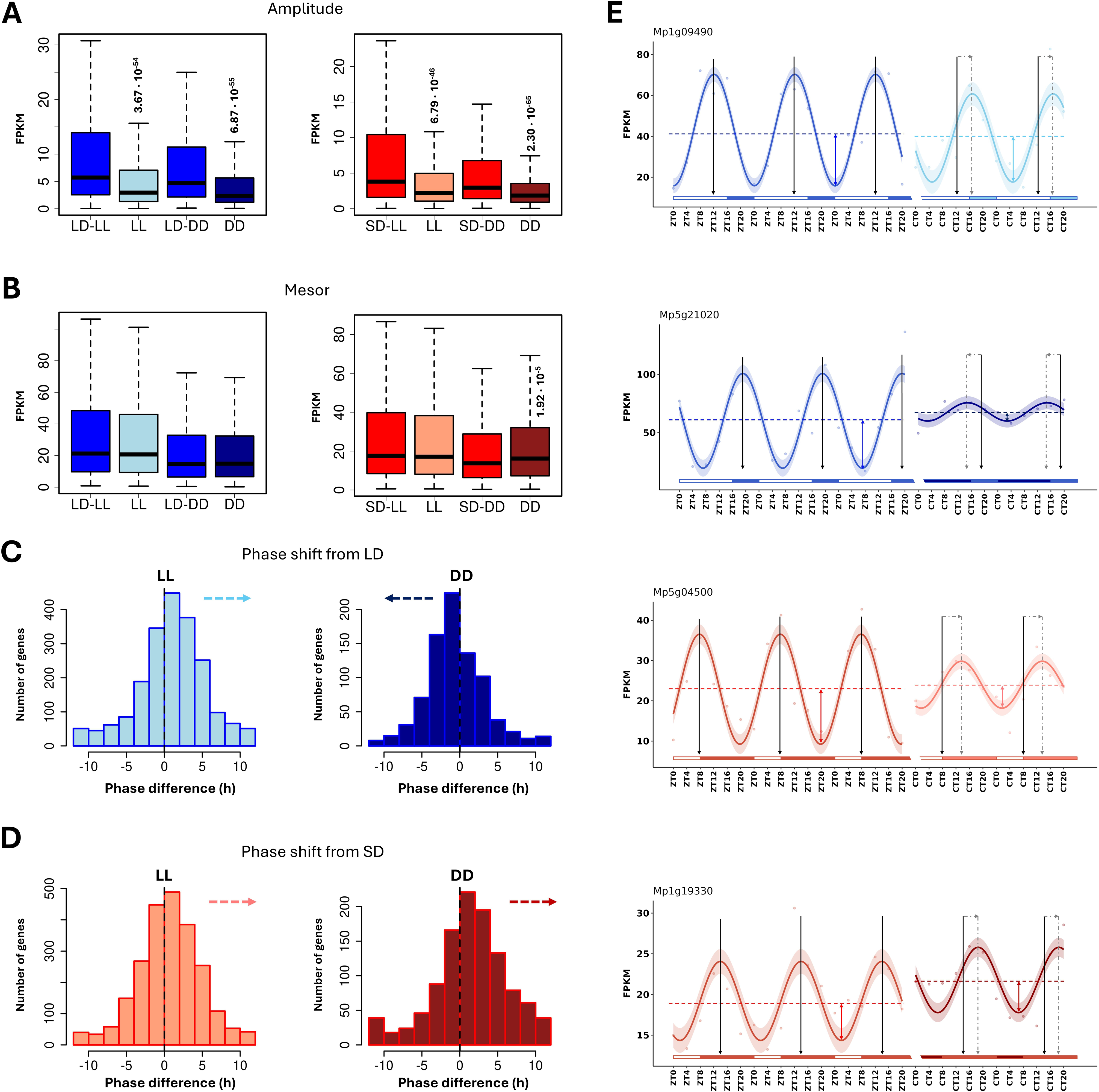
Gene expression profiles modifications after transitions to free-running conditions. **A)** Boxplots representing amplitude of different gene sets. Left panel: LD-entrained-LL-rhythmic genes amplitude under light/dark cycles (LD-LL) and constant light (LL), LD-entrained-DD-rhythmic genes amplitude under light/dark cycles (LD-DD) and constant darkness (DD). Right panel: SD-entrained-LL-rhythmic genes amplitude under light/dark cycles (SD-LL) and constant light (LL), SD-entrained-DD-rhythmic genes amplitude under light/dark cycles (SD-DD) and constant darkness (DD). Significance was computed according to Mann–Whitney–Wilcoxon nonparametric test using diel and free-running data. **B)** Boxplots representing mesor of different gene sets. Same sets as in Fig. 3A. Left panel: under LD. Right panel: under SD. Significance was computed as in the previous case. **C)** Phase shifts when plants are transferred from LD to LL (left) or DD (right). Dashed lines represent 0h-change. **D)** Phase shifts when plants are transferred from SD to LL (left) or DD (right). Dashed black lines represent 0h-change. **E)** Expression profiles of genes showing the differences in amplitude, mesor and phase mentioned in previous panels in transitions to free-running conditions. From top to bottom: transition from LD to LL, from LD to DD, from SD to LL and from SD to DD. Oscillating line corresponds to best-fitted cosinuidal expression curve, shaded area indicates amplitude error estimates and points represent expression values recorded at each time point. Double-headed arrows represent amplitude under every condition. Horizontal dashed lines represent mesor. Vertical black arrows mark phases under light/dark cycles, vertical dashed gray arrows mark LL and DD phases and horizontal dashed gray arrows represent delayed or advanced phases. White rectangles represent light periods, blue- or red-filled rectangles correspond to dark periods under LD or SD, respectively. Light blue rectangles correspond to subjective nights under LL, dark blue rectangles to subjective days under DD, light red rectangles to subjective nights under LL and dark red ones to subjective days under DD. ZTN, Zeitgeber time N, marks the time point N hours after dawn (lights on, ZT0). CTN, circadian time N denotes the time point N hours after subjective dawn. A discontinuity is shown on the time axis to indicate that samples were collected after 24 h acclimation to the corresponding free-running conditions.

Free-running conditions also affected the timing of maximum gene expression (phase). Under both photoperiods, transfer to LL conditions caused a general phase delay (**Fig. 3C-D, left panels)**. In constrast,transfer to DD conditions resulted in phase advances in LD-entrained plants (**Fig. 3C**), whereas SD-entrained plants showed similar phase delays to that observed under LL conditions (**Fig. 3D**). Representative examples of genes displaying these parameter shifts following transitions to free-running conditions are shown in **Fig. 3E**.

### Distinct dusk-responsive and photoperiod-specific circadian gene components modulate the global distribution of gene phases in diel cycles

The distribution of the number of genes that present their phase at each specific time point provides information regarding diel gene programs in different species, revealing shared and species-specific traits (Ferrari et al. 2019). In this respect, Marchantia has a predominantly dusk/night-phased transcriptome with rhythmic genes peaking preferentially around dusk under both LD and SD conditions, ZT16 under LD and ZT8 under SD (**Fig. 4A,E, left panels**). Marchantia gene phase distribution under LD resembles those shown under 12h light:12h darkness cycles by species such *as Chlamydomonas reindhardtii*, *Klebsormidium nitens*, *Physcomitrium patens*, and *Selaginella moellendorffii* (Ferrari et al. 2019). Nevertheless, the gene phase distributions differ in shape between photoperiods with LD being dominated by the dusk accumulation, whereas SD presented a bimodal distribution showing an additional nocturnal accumulation around ZT16.

**Figure 4.**
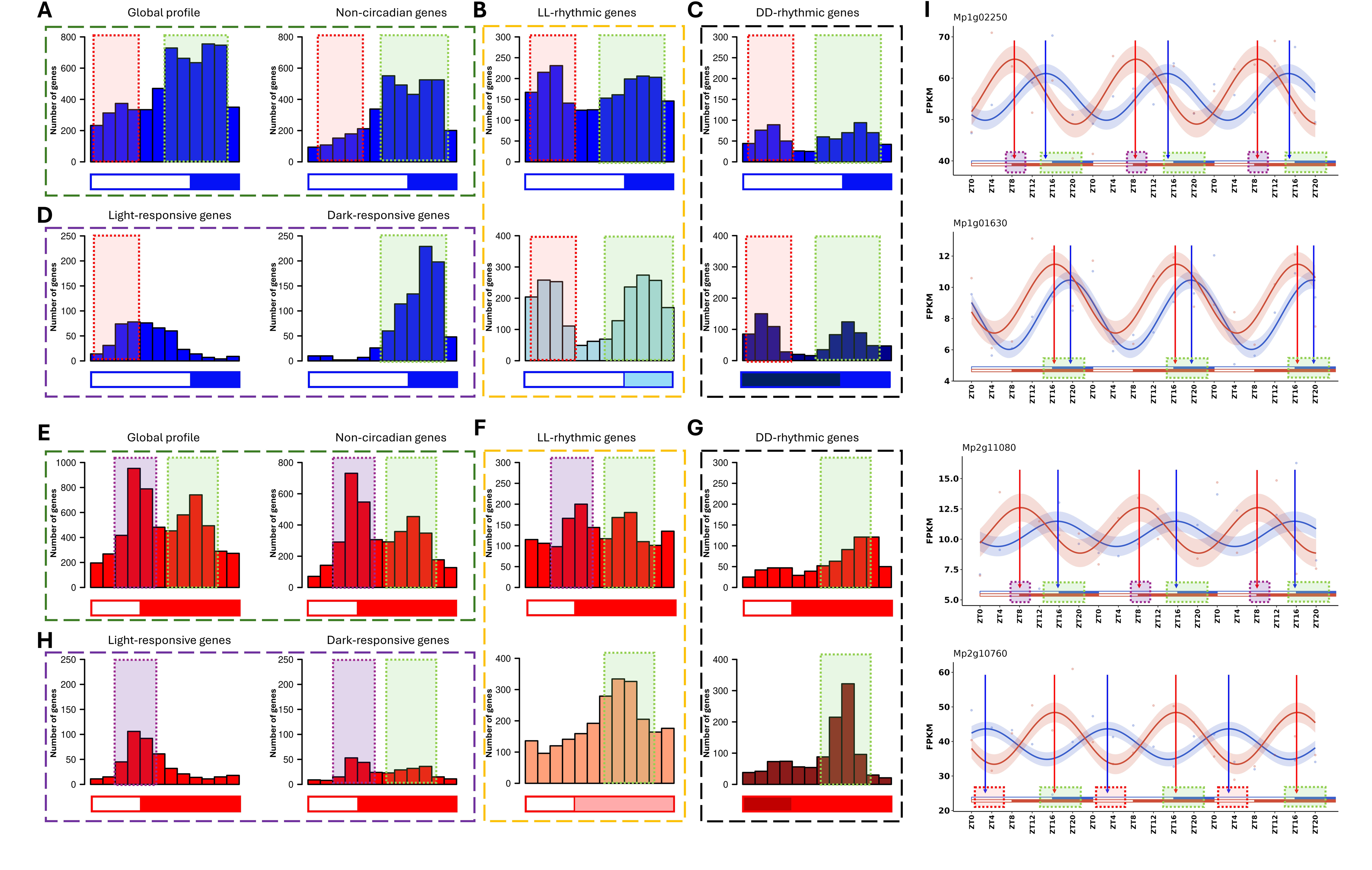
Global phase distributions and contributions from light/dark-responsive and circadian genes. Dotted red and green boxes indicate different phase accumulations and are included for reference. Color is consistent and indicates the same accumulation through different conditions under the same photoperiod. **A)** Phase distributions of rhythmic (left) and non-circadian-rhythmic (right) genes under LD-light/dark cycles. Green box corresponds to apparent phase accumulation at dusk, while red box indicates the time window at which a phase accumulation appears in constant conditions in the morning. Same for panels *B-D*. **B)** Phase distributions of LL-rhythmic genes under LD-light/dark cycles (top) and LL (bottom) after LD entrainment. **C)** Phase distributions of DD-rhythmic genes under LD-light/dark cycles (top) and DD (bottom) after LD entrainment. **D)** Phase distributions of light-responsive (left) and dark-responsive (right) genes under LD-light/dark cycles. Purple box corresponds to apparent phase accumulation at dusk, while red box corresponds to phase accumulation at ZT16. Same for panels *E-G*. **E)** Phase distributions of rhythmic (left) and non-circadian-rhythmic (right) genes under SD-light/dark cycles. **F)** Phase distributions of LL-rhythmic genes under SD-light/dark cycles (top) and LL (bottom) after SD entrainment. **G)** Phase distributions of DD-rhythmic genes under SD-light/dark cycles (top) and DD (bottom) after SD entrainment. **H)** Phase distributions of light-responsive (left) and dark-responsive (right) genes under SD-light/dark cycles. **I)** Expression profiles of illustrative genes showing the different contributions to phase accumulation depending on the photoperiod. From top to bottom: dark-responsive gene contributing to ZT8 accumulation under SD and ZT16 under LD, dark-responsive gene contributing to ZT16 accumulation under both SD and LD, light-responsive gene contributing to ZT8 accumulation under SD and ZT16 accumulation under LD, LL-rhythmic gene contributing to ZT4 accumulation under LD and ZT16 accumulation under SD. The corresponding phase accumulations are indicated by dotted boxes in the axis, whose colors correspond to those of panels *A-G*.

However, SD condition presented bimodality, with a second accumulation around ZT16 (green box) (**Fig. 4E, left panel)**, as that of LD. Bimodality has also been described for 12h light:12h darkness cycles in the liverwort (Tan et al. 2024) and other plants such as *A. thaliana* (Ferrari et al. 2019). Study of cycle–dependent rhythmic genes, i.e. rhythmic genes that did not exhibit cyclic behavior under LL or DD, revealed a distribution similar to the global one. This was expected, since they correspond to a large percentage of the total, but an underrepresentation of genes that do not align their phase with the dusk phase accumulation was also observed (**Fig. 4A,E, right panels**). This was the case for morning-phased genes under LD (red box) or the reduction in the night accumulation observed under SD (green box) compared to the total number of genes.

These global trends of gene phases result from the combination of responses to light and darkness with the circadian clock output (Mehta et al. 2021), and the different rhythmic gene sets could present different phase distribution depending on the light regime. Regarding free-running rhythmic genes expression under diel cycles, two phase accumulations were observed under LD, for both LL- and DD-rhythmic genes (**Fig. 4B-C, upper panels**). One of these peaks occurred just after the end of the light period (green box), while a new accumulation appeared during the morning (red box), presenting bimodality. Strikingly, under SD condition, despite its bimodality in the overall distribution of gene phases, only one accumulation was observed under light/dark cycles for DD-rhythmic genes (**Fig. 4G, upper panel**), coinciding with the nocturnal accumulation. Under LL, the double peak appeared, but with a diminished prominence compared with the overall distribution (**Fig. 4F, upper panel**). To find the origin of these photoperiod-dependent changes in modality, we studied the phase distribution under free-running conditions. Following LD entrainment, gene phase accumulations were observed at subjective morning and at subjective sunset under both LL and DD, coinciding with the phase accumulations presented by these gene sets under diel cycles (**Fig. 4B-C, lower panels**). In line with the phase shift results in transitions to constant conditions, both accumulations showed a delayed phase under LL and an advanced one under DD. In the case of SD, the morning accumulation disappeared completely, giving rise to unimodal distributions centered on the subjective night, corresponding to the second phase accumulation observed in diel cycles for this photoperiod (**Fig. 4F-G, lower panels**). Under LL, it can be observed that phase delays favored the disappearance of the first accumulation (**Fig. 4F**). Interestingly, although a generalized phase delay was also observed in SD-DD, the resulting distribution favored a sharp phase accumulation in the subjective night, earlier than that observed under diel cycles (**Fig. 4G**), similarly to LD. Therefore, the phase delay or advancement at the level of specific genes could result in the same global phase distribution under free-running conditions.

Influence of light and darkness on the cycle–dependent rhythmic genes response was analyzed separately. Thus, a light-responsive gene is one that, without maintaining its rhythmicity, progressively increases its expression over time under LL and decreases it under DD, and vice versa for dark-responsive genes. Under LD condition, a substantially higher number of dark-responsive genes (974) with respect to light-responsive ones (612) was observed. Their phase distributions showed how dark-responsive genes exerted a major influence on the appearance of the dusk accumulation, while phases of light-responsive genes were more evenly distributed during morning and noon, but not immediately after sunrise (**Fig. 4D**), contrary to recent findings in *A. thaliana* (Balcerowicz et al. 2021). Under SD condition, a greater number of light-responsive genes (769) were identified in comparison with dark-responsive genes (481), in contrast to LD. Of these, 144 dark-responsive and 221 light-responsive genes coincided with those under LD, around 30% of the total (Sup. Fig. S4A). No gene showed inverse regulation between photoperiods (Sup. Fig. S4B). Light-responsive genes again showed a delay in their phases relative to the start of the light period, thus contributing significantly to the dusk phase accumulation (**Fig. 4H, left panel**). Dark-responsive genes, on the other hand, exhibited rapid activation after the end of the light period, as under LD, and remained active for most of the long darkness period, decaying at the end (**Fig. 4H, right panel**). Therefore, they seemed to contribute to the two phase accumulations under SD. Hence, slow activation of light-responsive genes combined with rapid activation of dark-responsive ones gave rise to marked phase accumulations around dusk in both photoperiods, favoring the response to light under SD and to darkness under LD. On the other hand, slow activation of light-responsive genes in LD placed their expression outside the morning accumulation presented by free-running-rhythmic genes. This could explain the absence of that accumulation under diel LD condition, while the decay by the end of the night on the dark-responsive genes’ expression under diel SD emphasized the nocturnal accumulation present in all conditions. Examples of genes contributing to the described global accumulations are presented in **Figure 4I**.

Finally, these conclusions were contrasted with the phase distribution for bona fide-circadian genes which, being rhythmic in all conditions, allowed for the analysis of the circadian component in diel rhythms without the masking produced by cycle–dependent rhythmic genes. Under LD, the double accumulation detected under free-running conditions was also observed under diel cycles, with virtually no phase accumulation at noon and presenting the aforementioned -dynamics of DD-advancement and LL-delay (**Fig. 5A**). Under SD, the maximum accumulation was observed at night and around dawn in all conditions (**Fig. 5B**). The generalized DD phase advancement correlated with the separation of gene phases between a delayed morning accumulation and the night accumulation observed in the other conditions, retaining the ability to anticipate the light period under diel SD cycles.

**Figure 5.**
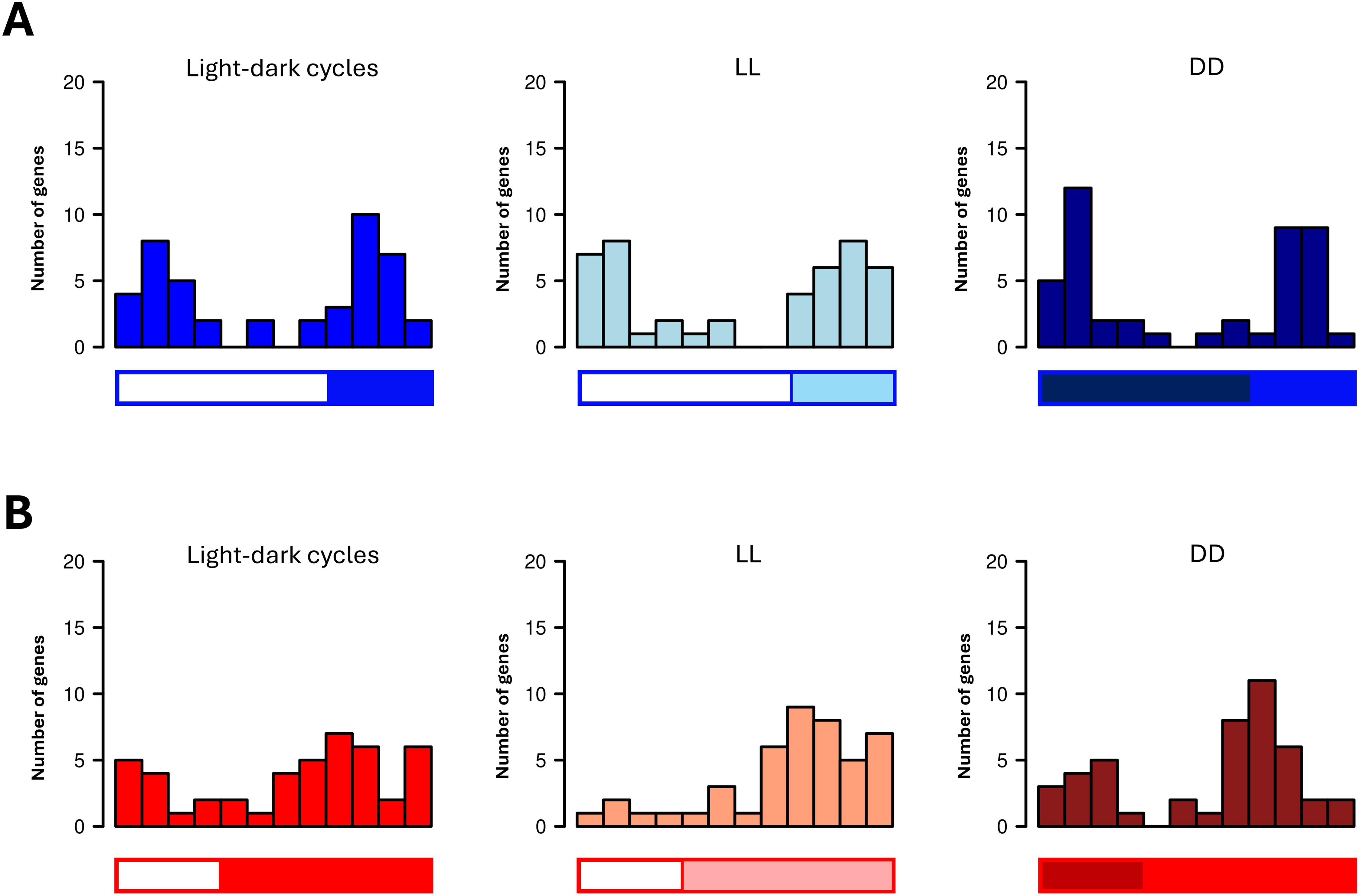
Phase distribution of bona fide-circadian genes **A)** under LD light/dark cycles (left), LL (center) and DD (right) after LD entrainment; **B)** under SD light/dark cycles (left), LL (center) and DD (right) after SD entrainment.

### Photoperiod-dependent changes in rhythmic gene expression profiles

Photoperiod was also found to affect wave parameters under light/dark cycles. A slight but significant increase in overall amplitude was found under LD condition, when compared with SD (p-value 0.02), despite the higher proportion of rhythmic genes in the latter (**Fig. 6A**, Sup. Fig. S5A). On the contrary, global mesor of rhythmic genes was higher for SD-entrained plants (p-value < 10^-5^) (**Fig. 6A**, Sup. Fig. S5B). Overall phase of rhythmic genes was anticipated under SD condition, exhibiting a significant difference of medians of 5.75h (**Fig. 6A-B**). Given the gene expression modality difference between photoperiods (bi-in SD, mono-in LD), this offset was also quantified gene-per-gene, resulting in a median difference of 3.54 h for genes showing rhythmicity under both photoperiods.

**Figure 6.**
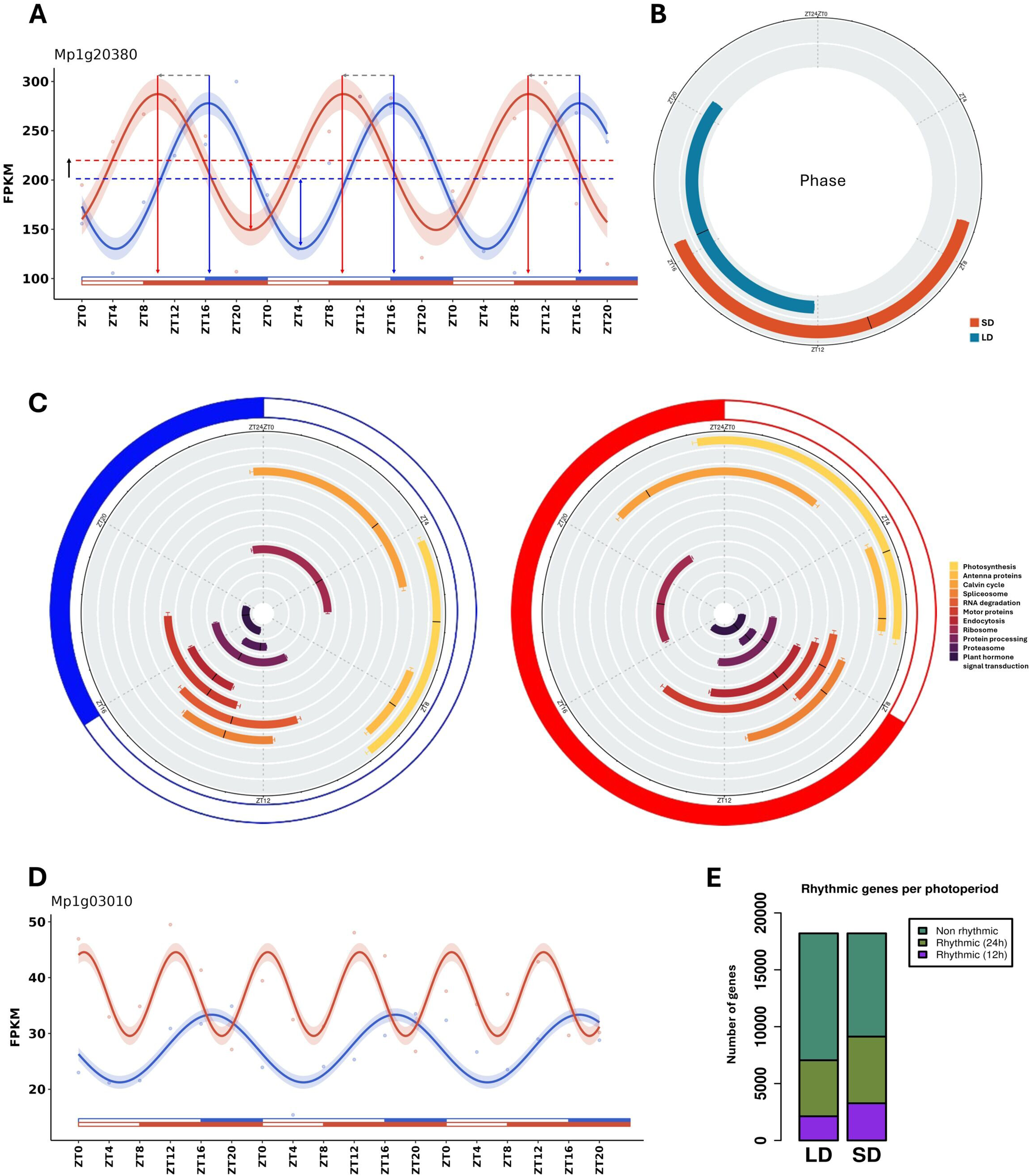
Photoperiodic effect on diel gene expression and biological programs. **A)** Representative expression profile of a gene example for the observed comparations between LD (blue) and SD (red) in the previous panels. Horizontal dashed lines indicate wave mesor, vertical arrows indicate phases, horizontal dashed gray arrows show photoperiodic phase shifts and double-headed arrows indicate wave amplitude. **B)** Circular boxplots representing phase of LD-(blue) and SD-rhythmic (red) genes. Whiskers indicate Standard Error of the Mean (SEM). **C)** Circular boxplots representing the distribution of phases of genes related to specific KEGG pathways. Left: under LD. Right: under SD. Whiskers indicate SEM. **D)** Representative expression profile of a gene example that presents only one daily peak under LD (blue) but two under SD (red). **E)** Stacked bar plots showing the number of arrhythmic, 24h-period-rhythmic and 12h-period-rhythmic genes under LD and SD. Colors are indicated in the legend.

Previous studies have linked photoperiod to differences in the diel timing of biological processes through changes in gene expression (Wu et al. 2023; do Nascimento et al. 2025; Romero-Losada et al. 2025). While many processes exhibit diel rhythms, with gene expression confined to specific temporal windows, others are independent of light–dark cycles and display uniform distribution of their phases across the day/night (Panchy et al. 2014). Among diel-clustered processes, a further distinction exists between those following the global phase distribution and those deviating from it. The former likely reflects temporal coordination driven by periods of increased DNA accessibility for transcription, such as during dark spans (Pittendrigh 1993). In contrast, the latter group displays specific and tightly scheduled gene expression, suggesting that precise temporal control is likely functionally important for these processes (Zhang et al. 2016). For instance, this could be the case for processes that rely on light. Distinct diel timings of gene phases for specific biological processes were identified by performing circular cohesion analysis over GO terms-associated genes under both photoperiods. 1279 GO terms presented a non-random unimodal distribution of gene phases under LD and 1212 under SD conditions (Sup. Table S3-S4). Identity of these GO terms were largely conserved between photoperiods, with only 346 LD-specific and 280 SD-specific ones (Sup. Fig. S5C) While the expression of most GO terms-associated genes mimicked that of the global gene phase distribution, 451 under LD and 256 under SD exhibited a significant distinct organization of associated gene phases, suggesting high importance of diel timing in their function (Sup. Fig. S5D). Among those that presented an overall GO-term phase distribution (i.e. the phase distribution of genes sharing a specific GO term) different from the global one, those related to transcription regulation and chloroplast-associated processes, photosynthesis, cytoskeleton organization, organelle assembly, translation, ribosome biogenesis and protein catabolism stood out (Sup. Table S5-S6). We detected photoperiodic changes in GO-term phase distribution agreeing with the advanced gene phases observed under SD. 572 GO-terms significantly advanced their phases under SD, and only “protein-DNA complex” delayed it (Sup. Table S7-S9).

As for specific pathways, the same analysis was performed based on KEGG pathways instead of GO terms. Non-uniformity was identified for 73 pathways under LD and 69 under SD, 55 of which were shared between conditions (Sup. Fig. S5E) (Sup. Table S10-S11). 25 and 21 pathways showed deviation from global phase distribution in LD and SD conditions, respectively, with 16 shared (Sup. Table S12-S13) (Sup. Fig. S5F). Photoperiodic rearrangement of gene phases for these specific pathways favored in all cases the anticipation in short photoperiod, highlighting the advancement of pathways such as photosynthesis, antenna proteins, carbon fixation by Calvin cycle, spliceosome, RNA degradation, motor proteins, endocytosis, ribosome, protein processing in endoplasmic reticulum, proteasome, and plant hormone signal transduction, among others (**Fig. 6C**) (Sup. Table S14).

Overall, most biological processes exhibited temporal coordination in their gene phases aligned with the phase distribution at whole-transcriptome level. However, key processes tightly clustered at specific time points not coinciding with the global phase accumulations and exhibited photoperiod-dependent timings. Among these, photosynthesis stood out, whose genes were expressed at the end of the dark period and the beginning of the light period under SD condition, while, under LD, it was delayed until after ZT4. In both cases, antenna proteins were expressed towards the middle of the light period, which may be related to processes responding to radiation and light stimuli, whose genes phases overlapped temporally. Calvin cycle genes also advanced their phases under SD, showing high expression already at dawn, unlike under LD. ROS-response genes were mainly expressed during the morning, suggesting that light stress was primarily responsible for their production. Ribosome assembly exhibited gene expression diametrically opposed to that of the proteasome, occurring towards the end of darkness under SD and at the beginning of light under LD, while protein catabolism was concentrated around ZT8 under SD and ZT14 under LD. Transcription-related genes peaked around dusk under both photoperiods, while their positive regulation occurred two hours earlier. Organelle organization and cytoskeleton remodeling also occurred earlier under SD. DNA repair was mostly limited to the dark period, at ZT20 under LD and at ZT15 under SD. This temporal sequence in key biological processes broadly agrees with that determined for other plant species, like *Chlamydomonas reindhardtii* (Panchy et al. 2014).

The study of the period of rhythmic genes showed that the global bimodality detected under SD diel cycles correlated with an increase in the number of genes exhibiting a double peak of diel expression with respect to LD (3267 vs 2125) (**Fig. 6D-E**). 722 of the genes showing this double daily peak under SD presented a single peak under LD, while only 451 displayed the opposite behavior. Unlike what has been observed in the microalgae *Ostreococcus tauri* (Romero-Losada et al. 2025), these photoperiod-specific double peaks did not seem to be associated with the overlap of distinct out-of-phase photo- and skotoperiodic waves, which would result in single and alternating peaks in LL and DD conditions, not observed in our data.

### Specific transcription factor binding sites are associated with circadian and light/dark-responsive gene expression

One of the most important regulatory loops in maintaining diel rhythms is exerted by transcription factors (TFs) that regulate the rhythmic expression of genes trough their binding to specific *cis* elements in gene promoters (Petersen et al. 2022; Wu et al. 2023). To determine potential regulators giving rise to the diel organization of biological processes described, we analyzed the relationship between Transcription Factor Binding Sites (TFBS) in gene promoters and their phases. Specific-TFBS-associated gene sets that deviated from the overall expression pattern comprised 27 motifs under LD but only 11 under SD condition (Sup. Tables S15–S16). These TFBS should therefore be involved in clustering gene expression at specific times of the day that differ from the general phase distribution of rhythmic genes. Ten of them played this role under both photoperiods: abf1, ABF2, ABI5, ANAC057, At2g45680, bZIP28, bZIP48, E-box, GBF5, and HSF3. These results suggested an important role for basic leucine zipper (bZIP) motifs in determining specific timings of gene expression, as in other plants (Viana et al. 2021). Analysis of the expression of their associated genes revealed a delayed distribution with respect to the global phase accumulations (Sup. Fig. S6A). To avoid confounding effects derived from multiple TFBS per promoter, we repeated the analysis using genes with few co-occurring motifs, observing phase accumulations near dawn under LD and at ZT20 under SD (Sup. Fig. S6B). Thus, the phase for genes exhibiting bZIP motifs in their promoters could be associated with gene expression towards the beginning of the light period.

A TFBS enrichment analysis was also performed for genes that showed maximum expression centered on ZT0/4/8/12/16/20 under each photoperiod (**Fig. 7A**). We detected motifs that appeared at earlier time points under SD compared to LD, such as G-box at ZT8 vs ZT4 or WRKY at ZT16 vs ZT12. Significant contribution of G-box was observed in the early hours of the day and of Teosinte branched 1/Cycloidea/Proliferating cell factor (TCP)-binding motifs during the light period, giving way to those recognized by WRKY, myeloblastosis (MYB), and APETALA2/Ethylene-Responsive Element Binding Protein (AP2EREBP) TFs during the dark period under both LD and SD. bZIP and TCP motifs, motifs known to play a central role in circadian regulation in *A. thaliana* (Wu et al. 2016; Viana et al. 2021), were predominant in promoters of photosynthetic, chloroplast organization and ribosome assembly genes, focusing their expression at the beginning of the light period. Conversely, genes related to transcription regulation, proteolysis, and signaling were mostly expressed in the night period, exhibiting high abundance of NAM/ATAF1/CUC2 (NAC) and WRKY motifs in their promoters.

**Figure 7.**
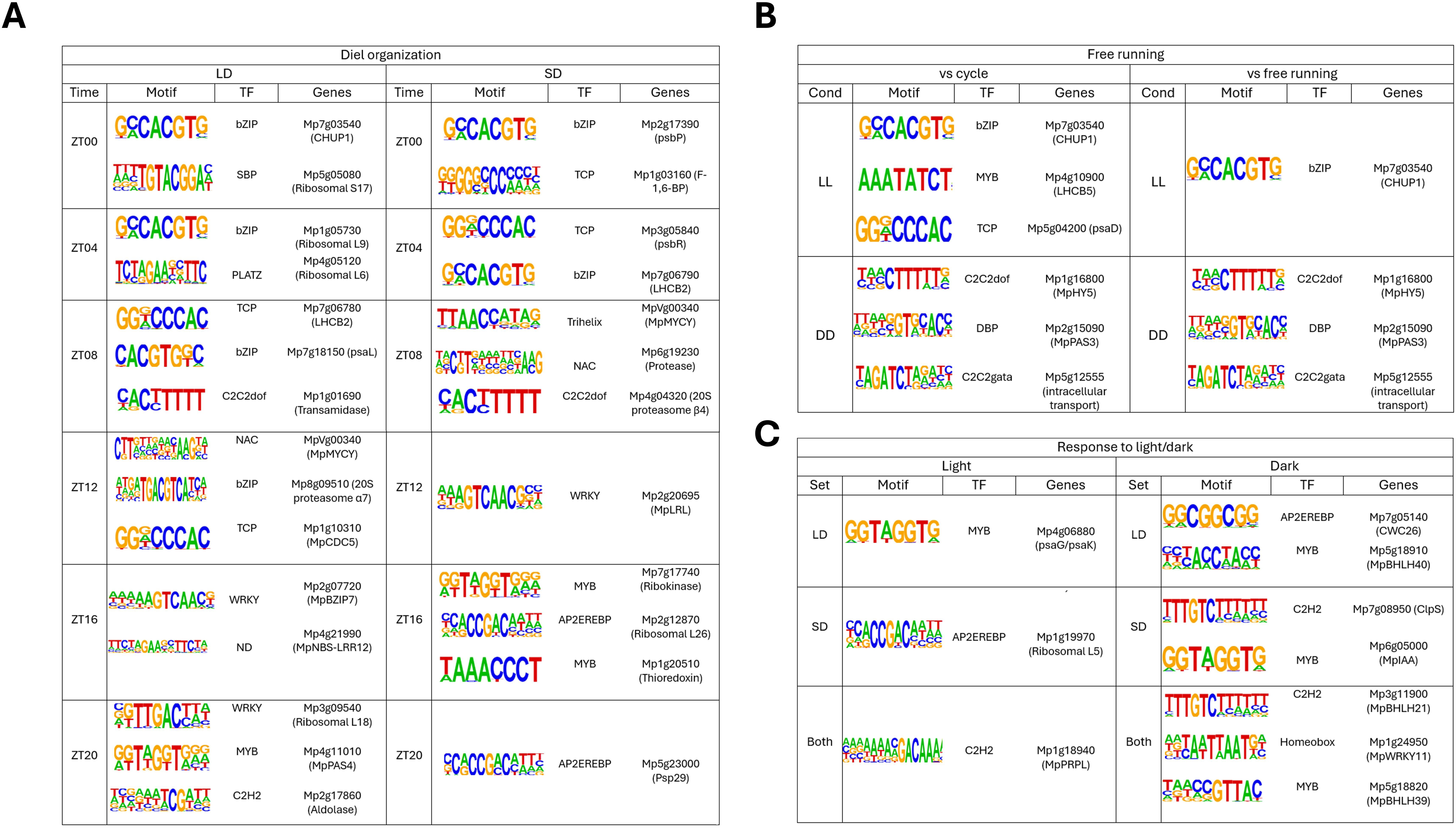
TFBS enrichment in different gene sets showing enriched motif, TF family that recognizes it and gene examples presenting it in their promoters. **A)** Enrichment for each time point gene cluster for LD-(left) and SD-rhythmic genes (right) when compared with the complete LD- or SD-rhythmic-genes sets, respectively. **B)** Enrichment for LL-(top) and DD-rhythmic genes (bottom) when compared with the complete rhythmic-genes set (left) or with the free-running-rhythmic-genes set (right). **C)** Enrichment for light-(left) and dark-responsive genes (right) in LD (top), SD (mid) or both (bottom) when compared with the complete LD-rhythmic-genes set, SD-rhythmic-genes set and the intersection of both, respectively.

Regarding circadian regulation, TFBS enrichment of LL-rhythmic genes compared to diel cycles-rhythmic genes revealed motifs recognized by bZIP (G-box), MYB-related (Evening Element), and TCP TFs (**Fig. 7B**), consistent with circadian clock targets in other plants (Lau et al. 2011; Wu et al. 2016; Viana et al. 2021). Meanwhile, DD-rhythmic genes presented DNA-binding with one finger (C2C2dof), GATA-binding (C2C2gata) and DBP motifs in their promoters. A separate question was whether there existed TFBS enrichment for LL- or DD-rhythmic genes compared to rhythmic genes in any free-running condition. In this case, LL-rhythmic genes showed enrichment in G-box motifs recognized by bZIP and bHLH (such as those recognized by *PIF* TFs) TFs, while DD-rhythmic genes maintained the same results (**Fig. 7B**). Hence, TCP and EE may support rhythmicity in LL but were less specific to that condition than the G-box, while C2C2dof, C2C2gata, and DBP were strongly favored in DD-rhythmic-genes promoters. Moreover, only one weakly enriched MYB-related motif was detected when comparing free-running-rhythmic genes against diel-cycle-rhythmic ones, suggesting that rhythmicity in LL and DD did not stem from a shared regulation.

As detailed before, the main non-circadian contribution to diel gene expression is that of light- and dark-responsive genes. Under LD, we detected significant enrichment of MYB motifs in light-responsive-genes promoters, while dark response was associated with AP2EREBP and MYB motifs (**Fig. 7C**). Under SD, this changed to AP2EREBP for light-responsive genes and C2H2 and MYB for dark-responsive ones (**Fig. 7C**). The results implied that there existed no significant conservation of TF regulation between light and darkness responses under different photoperiods. Genes consistently classified as either light- or dark-responsive in both photoperiods were also analyzed, revealing the existence of a C2H2 motif enriched in the light response under both LD and SD, also present in the response to darkness along with Homeobox and MYB ones (**Fig. 7C**).

When applying the same analysis to elucidate photoperiod-specific rhythmicity maintenance in transitions to free-running, no enriched motifs appeared. Therefore, it was not expected that there existed a TF that helped to maintain the rhythmic character of genes preferentially from one or another photoperiod, and the differences in the percentages of rhythmic genes (e.g., in SD-DD) were likely due to the pre-transition amplitude of gene expression. Finally, genes with double peaks under SD showed enrichment in bZIP motifs (bZIP3 and GBF5), while, under LD, this phenomenon could not be attributed to any specific motif (Sup. Fig. S6C).

In summary, combined results from circular distribution of gene expression related to specific TFBS and enrichments in different gene sets suggested involvement of the G-box motif in the circadian organization of genes and gene expression at dawn but not related to light-responsive genes. During dusk and night, C2H2, MYB, and AP2EREBP motifs, related to the response to darkness, appeared to be more important.

### Regulation of diel and circadian rhythms by transcription factors under different photoperiods

Candidate genes for the central circadian clock of Marchantia have been proposed by (Linde et al. 2017) and (Lagercrantz et al. 2021), based on comparisons with *A. thaliana* clock and the maintenance of their expression levels under ND-LL. However, their expression patterns under different photoperiods or constant darkness have not been yet systematically explored. In **Table 1**, we summarize the expression rhythms of these genes in the transitions from both photoperiods to LL and DD and their comparison with the results of (Lagercrantz et al. 2021). Reported results for ND-LL broadly coincided with those obtained in this study for LD-LL and SD-LL, with the exceptions of *WD REPEAT-CONTAINING PROTEIN 2* (*MpWDR2*), non-rhythmic under LD-LL, and *MpTOC1*, whose weak rhythms determined in (Lagercrantz et al. 2021) did not reach significance in any of the most extreme photoperiods. Drastic differences were observed under DD condition, with a generalized reduction in rhythmicity, especially after SD-entrainment, under which only *CONSTANS-LIKE 2* (*MpCOL2*) maintained significant rhythmicity. Under LD-DD, *MpCOL6* and *MpDET1* maintained their cyclic profile.

**Table 1.** Rhythmic character of *Marchantia polymorpha* candidate circadian clock genes. Adjusted p-values of rhythmicity assessment in each condition are provided, along with those reported in (Lagercrantz et al. 2021) for ND-LL.

| Symbol | ID | LD-LL | SD-LL | LD-DD | SD-DD | ND-LL* |
| --- | --- | --- | --- | --- | --- | --- |
| **MpCOL6** | Mp5g05940 | 2.47E-04 | 3.31E-04 | 2.07E-02 | 3.70E-01 | 6.69E-07 |
| **MpPRR** | Mp2g16560 | 4.74E-04 | 5.56E-04 | 1.68E-01 | 3.82E-01 | 9.27E-06 |
| **MpCDF1** | Mp1g18540 | 1.16E-03 | 7.11E-03 | 1.07E-01 | 3.43E-01 | 5.20E-05 |
| **MpLNK** | Mp1g10620 | 2.11E-03 | 7.08E-04 | 6.95E-02 | 3.85E-01 | 9.35E-07 |
| **MpDET1** | Mp8g00970 | 4.49E-03 | 6.65E-04 | 3.32E-02 | 1.99E-01 | 3.13E-07 |
| **MpCOL2** | Mp2g16350 | 1.24E-02 | 9.92E-03 | 1.01E-01 | 2.98E-02 | 6.52E-05 |
| **MpRVE** | Mp2g14310 | 3.67E-02 | 1.31E-02 | 6.34E-02 | 2.99E-01 | 6.13E-02 |
| **MpWDR2** | Mp5g01830 | 2.07E-01 | 4.59E-03 | 1.05E-01 | 2.82E-01 | 9.83E-04 |
| **MpTOC1** | Mp3g09460 | 3.32E-01 | 1.75E-01 | 2.60E-01 | 5.37E-02 | 1.57E-02 |
| **MpGI** | Mp1g13750 | 3.73E-01 | 2.41E-02 | 2.25E-01 | 3.45E-01 | 1.74E-02 |
| **MpELF3** | Mp1g10870 | 4.03E-01 | 8.39E-02 | 1.43E-01 | 5.05E-02 | 1 |

In addition to the previously proposed genes, we explored TFs exhibiting cyclic behavior in all transitions to free-running conditions to select new potential candidates with a possible association with the circadian clock of Marchantia. Under LL, 71 were rhythmic when entrained under SD, 33 under LD, and 18 under any photoperiod (Sup. Fig. S7A). Gene phases of the last group clustered at the beginning of the light period under LD-LL and at ZT16/20 under SD-LL (**Fig. 8A**). Under DD, these numbers were reduced to 34, 26 and 7, respectively (Sup. Fig. S7B). DD-rhythmic genes presented their phases strongly centered around ZT16 under SD-DD, while those under LD-DD clustered more loosely around sunrise (**Fig. 8B**). TFs expression pattern under constant conditions therefore correlated with the bimodality of gene expression detected in transitions from LD and the ZT16 peak from SD. Only three TFs showed significant rhythms under all conditions: *ELONGATED HYPOCOTHYL 5* (*MpHY5*, *Mp1g16800*, **Fig. 8C**), *MpTRIHELIX31* (*Mp4g09730*, **Fig. 8D**) and *MpTRIHELIX11* (*Mp2g08360*, **Fig. 8E**). *HY5* is a bZIP TF shown to bind G-box motifs (Chen et al. 2013; Gangappa and Kumar 2017). The *MpHY5* transcript peaked at the end of the night under SD, LD, and all free-running conditions except LD-LL, under which it peaked at dawn. The two Trihelix TFs showed an expression pattern centered on dawn, suggesting that the three TFs could play a role in predicting the onset of the subjective light period under constant conditions.

**Figure 8.**
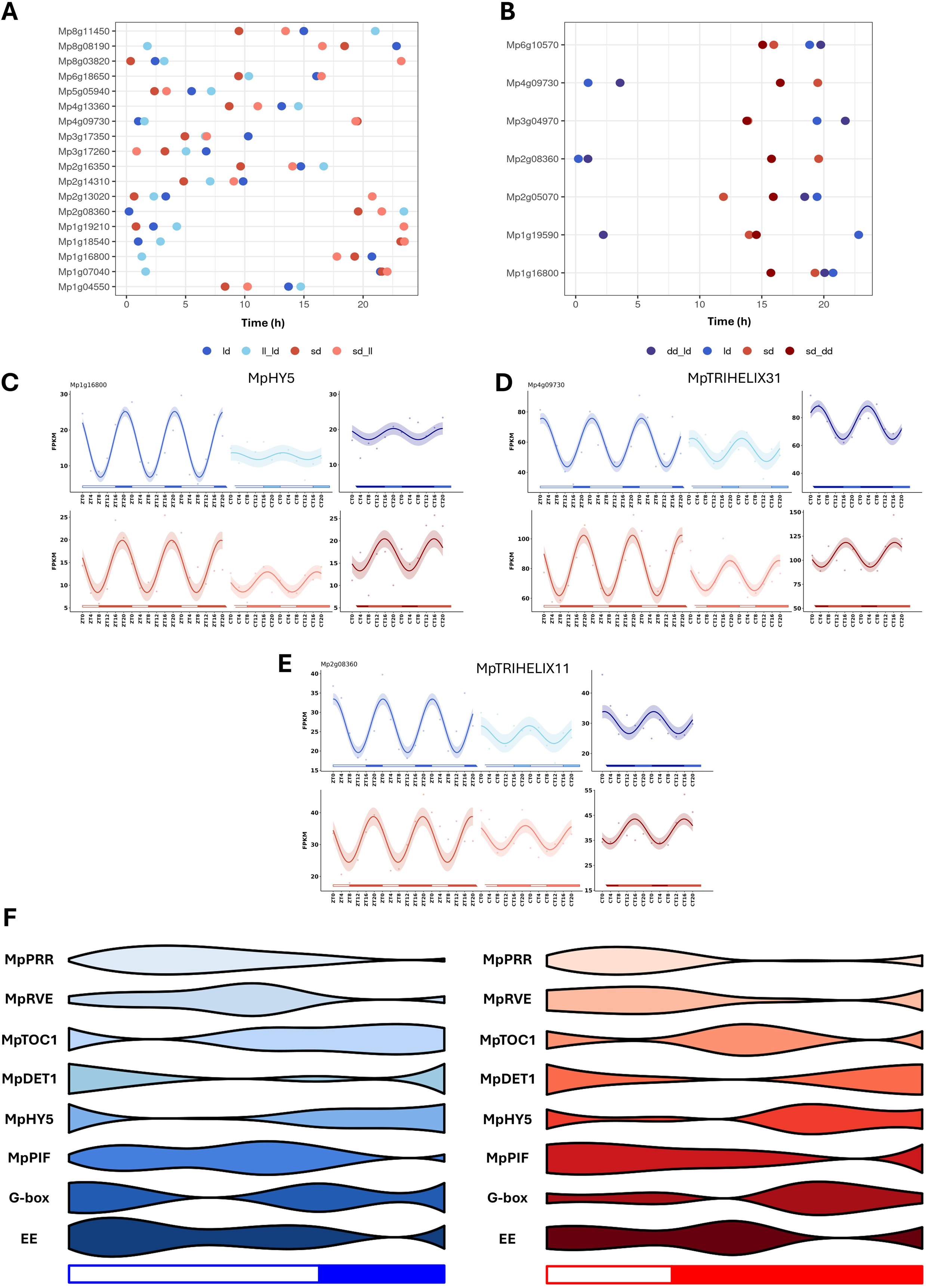
Photoperiodic differences in TF expression profiles. **A)** Phase of LL-rhythmic genes in both photoperiods. Light regime of each phase is indicated in the legend. **B)** Phase of DD-rhythmic genes in both photoperiods. Light regime of each phase is indicated in the legend. **C)** Expression profile of *MpHY5* under every condition. **D)** Expression profile of *MpTRIHELIX31* under every condition. **E)** Expression profile of *MpTRIHELIX11* under every condition. **F)** Violin plots representing expression profiles trough light/dark cycles of, from top to bottom: *MpPRR*, *MpRVE*, *MpTOC1*, *MpDET1*, *MpHY5*, *MpPIF*, G-box-containing and EE-containing genes (both are reduced sets filtered by genes exhibiting less than 20 simultaneous motifs in their promoters). Violin widths represent the corresponding average gene expression of three days of measurements. Left: LD. Right: SD.

Another protein whose impact on the circadian clock of Marchantia has recently been highlighted is *MpDET1* (Lagercrantz et al. 2021). *DET1* binds to *CCA1* and co-represses its targets in *A. thaliana* and is dependent on the expression rhythms of *CCA1* for its function (Lau et al. 2011). However, in Marchantia, it exhibits rhythmicity under ND-LL (Lagercrantz et al. 2021) with a phase close to dawn. In our study, cyclic expression was observed under all study conditions except SD-DD, anticipating dawn under diel cycles, with small phase shifts between photoperiods (Sup. Fig. S7C). A known function of this protein in *A. thaliana* is the degradation of *HY5* through its association with *COP1*, allowing the integration of light and temperature signals into the clock (Cañibano et al. 2021). In Marchantia, we observed how in both photoperiods, the expression pattern of *MpDET1* and *MpHY5* was consistent with a model in which *MpHY5* played a role in promoting gene expression at dawn and *MpDET1* acted as an inhibitor of its expression (**Fig. 8F**). Furthermore, these mRNA distributions correlated with the expression increased in genes containing a G-box in their promoter during the period of high *MpHY5* expression and its reduction shortly afterwards. While *HY5* acts as an expression activator of its target genes, *PIF* is a well-known TF that represses *HY5* targets in *A. thaliana*, regulating the expression of these genes through a temporal abundance pattern different from *HY5* (Chen et al. 2013). Marchantia possesses a single *MpPIF* (Chen et al. 2013), which showed rhythmicity under both photoperiods and transitions to LL, but not to DD. Its expression increased coinciding with the decreased levels of *MpHY5* transcript and genes that carry a G-box in their promoter (**Fig. 8F**), suggesting that it may have a similar function to that exhibited in *A. thaliana* regarding antagonistic regulation of *HY5*.

Expression of genes carrying the Evening Element (EE) in their promoter, a motif recognized by *CCA1* that represses the expression of its targets during the morning in *A. thaliana*, nevertheless showed an increase at the beginning of the day. Thus, the existence of *MpDET1* with a phase close to dawn did not seem to be sufficient to maintain the repression of these genes in the absence of *CCA1*, implying that this loop should have evolved divergently in liverworts and tracheophytes. Finally, M*pPRR*, *MpRVE*, and *MpTOC1* appeared to display sequential expression spanning the whole day under LD but did not show high expression in the late hours of the night under SD (**Fig. 8F**). Their expression pattern also supported conclusions from (Lagercrantz et al. 2020): *MpPRR* and *MpTOC1* mutually repress each other and repress *MpRVE*, which in turn promotes the expression of both. Interestingly, *MpTOC1* showed a second, smaller peak of expression at dawn under SD, decaying shortly afterwards until its activation at the beginning of the dark period.

### Short photoperiod-entrained plants exhibited anticipated diel profiles of Fv/Fm and NPQ

As mentioned above, the expression of photosynthetic genes undergoes significant adaptation to the seasonal photoperiod with respect to their phases in diel cycles, necessary to optimize energy conversion (Masojídek et al. 2021). The anticipation detected under SD condition should impact on the expression of some or all the four complexes involved in this process: PSII, b6f cytochrome, PSI, and ATP synthase. Individual analysis of the constituent genes determined a generalized anticipation in their phase under short photoperiod (**Fig. 9A-E**). Also, occurrence of double daily expression peaks under this condition was observed, generally resulting in one peak anticipating dawn and another occurring towards the middle of the light period (**Fig. 9A-E**). On the other hand, LD condition presented a single peak, broader than those of SD, extending over most of the light period. Specifically, profiles with 2 peaks in SD condition and only one in LD could be observed in genes such as *petM*, *atpF* and *LHCB6*, coding for key components of the complexes (**Fig. 9B-E**). Many of these genes maintained oscillations under LL but not DD, supporting the existence of more robust photosynthetic rhythms under the former. In turn, this might be associated with the lower Relative Amplitude Error (RAE) values found for LL in (Cuitun-Coronado et al. 2022). To analyze whether the transcriptional advancement also produced a physiological anticipation in photosynthetic activity, Pulse-Amplitude-Modulation (PAM) fluorometry measurements were performed to quantify the maximum quantum efficiency of PSII throughout diel cycles (Fv/Fm) (**Fig. 9F**). Diel fluctuations in this parameter exhibited a rhythmic pattern with a 24 h period in both photoperiods, peaking at ZT16 under LD and ZT12 under SD, correlating with the advancement of photosynthetic processes identified in the previous analysis, with a slightly lower value (4%) under SD compared to LD. In contrast, Fv/Fm reached its minimum 12 h later in both cases, also exhibiting a 6% reduction under the latter conditions.

**Figure 9.**
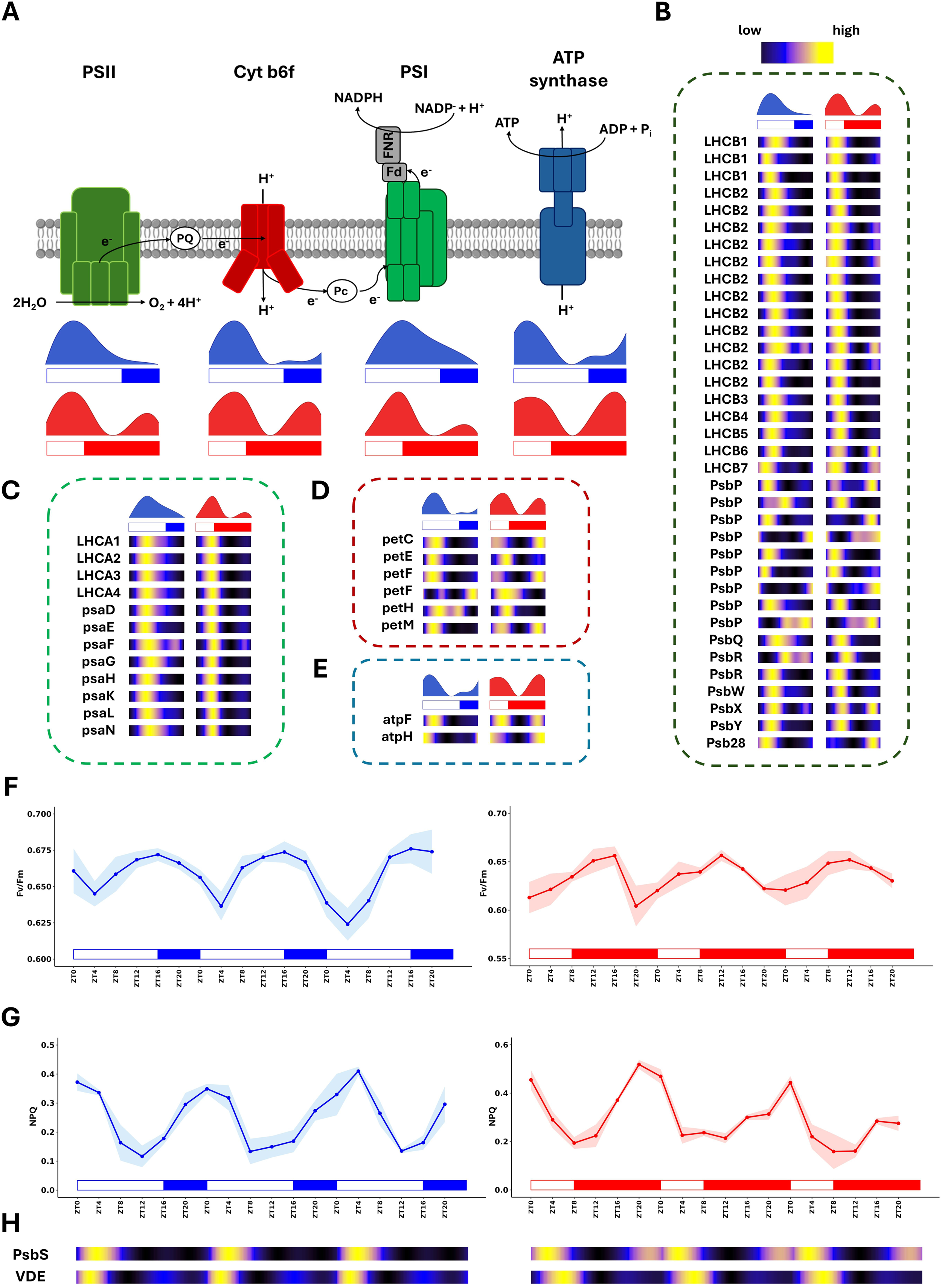
Photoperiodic effects on photosynthetic physiology and gene expression. **A)** Schematic representation of photosynthetic transport with the averaged expression profiles for all genes comprising PSII, cytochrome b6f (Cyt b6f), PSI, and ATP Synthase complexes under LD (blue) and SD (red) conditions. **B)** Heatmaps showing transcript abundance for PSII genes under LD (left, blue) and SD (right, red) conditions. Black represents low, blue medium, and yellow high abundances. Genes are indicated by their gene symbols. **C)** Heatmaps showing transcript abundance for PSI genes under LD (left, blue) and SD (right, red) conditions. **D)** Heatmaps showing transcript abundance for cytochrome b6f genes under LD (left, blue) and SD (right, red) conditions. **E)** Heatmaps showing transcript abundance for ATP synthase genes under LD (left, blue) and SD (right, red) conditions. **F)** Operational efficiency of PSII during 3 consecutive days under LD (left, blue) and SD (right, red) conditions. Shaded area corresponds to Standard Error of the Mean (SEM). **G)** NPQ values during 3 consecutive days under LD (left, blue) and SD (right, red) conditions. Shaded area corresponds to Standard Error of the Mean (SEM). **H)** Heatmaps showing transcript abundance for NPQ-related genes *PsbS* and *VDE* during 3 consecutive days under LD (left) and SD (right) conditions.

In our experimental design, we also identified a rhythmic behavior in diel cycles for non-chemical quenching (NPQ) component, peaking at ZT0 under SD and ZT4 under LD (**Fig. 9G**). Unlike Fv/Fm, the NPQ peak was narrower, reaching its minimum 8 hours later in both photoperiods. In general, NPQ values under SD were 17% higher than under LD. Therefore, NPQ seemed to be associated with a protective function during the transition from the dark to the light period and appeared to play a particularly important role in plants adapted to short photoperiod. Thus, under SD, the involved molecular mechanisms were anticipated to present the NPQ peak already at dawn, as well as maintaining values higher than those of LD throughout the day. As for the delay under LD, this allowed high NPQ values to be maintained during the hours of maximum light intensity of the extended lighting period, limiting photochemistry at these time points. Among the main mechanisms of NPQ are ΔpH across the thylakoid membrane and the activation of *PsbS* and *VIOLAXANTHIN DE-EPOXIDASE* (*VDE*) proteins (Guardini et al. 2025), whose associated genes showed high expression around dawn (**Fig. 9H**). mRNA levels of *PsbS* rose from ZT16 under SD, in contrast to LD, under which they did not increase until dawn. *VDE* presented a double peak in both photoperiods, one coinciding with the first hours of light and the second coinciding with the progressive nocturnal increase in NPQ, around ZT16. In addition, *PsbS* proved to be rhythmic under all constant conditions except SD-DD, while *VDE* presented weaker rhythms, maintaining them only under LD-LL.

Therefore, these results suggested that an early reorganization of the light harvesting complexes and anticipation in the expression of key genes took place under SD condition so that at dawn the photochemical capacity started to rise to take advantage of the short photoperiod. On the other hand, under LD it needed a previous adaptation period and presented its minimum value at ZT4. This phenomenon occurred in conjunction with the increase in NPQ levels around the beginning of the illumination period, presumably because the night-to-day transition defines a stress barrier around sunrise in Marchantia. In turn, this boundary reduced photosynthetic rate under all photoperiods to avoid photodamage of the photosynthetic machinery.

### Photoperiod-dependent chloroplast photorelocation and photosynthetic physiology to optimize light harvesting and photoprotection

In addition to Fv/Fm and NPQ, another major determinant of the amount of light energy that can be assimilated by the plant is the availability of chloroplast area exposed on the cell surface. Chloroplast photorelocation between anticlinal and periclinal cell walls has been described to be mediated by phototropin kinase signals in land plants in the presence of blue light through its cis-/trans-phosphorylation state (Kato et al. 2021; Noguchi et al. 2025), encoded in a single copy by the *MpPHOT* gene in Marchantia (Komatsu et al. 2014). Moreover, *CHLOROPLAST UNUSUAL POSITIONING 1 (MpCHUP1)*, *NRL PROTEIN FOR CHLOROPLAST MOVEMENT 1 (MpNCH1) KINESIN-LIKE PROTEIN FOR ACTIN-BASED CHLOROPLAST MOVEMENT (MpKAC*) and *RETICULON* (*MpRTN1*) genes have been identified as responsible for this movement in response to light in the liverwort (Suetsugu and Wada 2016; Suetsugu et al. 2016; Ishikawa et al. 2022; Yamamoto-Negi et al. 2024) in accordance with the proposed model (**Fig. 10A**). Rhythmic expression for all these genes was detected in both photoperiods showing advancement under SD with respect to LD (**Fig. 10A).** *MpCHUP1* and *MpKAC* showed a narrow expression peak around dawn in both conditions, possibly making the motility system available to perform its function during the light period. As for signaling, phototropin expression exhibited a clear anticipation under SD, increasing from ZT20, and 4 hours later under LD. This suggested transcriptional regulation of the process in addition to the known control by receptor phosphorylation. *MpNCH1* only mediates the accumulation response and was expressed at the end of the night under SD and at the beginning of the light period under LD, favoring a potential early accumulation of chloroplasts under shorter photoperiod. On the other hand, *MpRTN1*, mediator of avoidance response, peaked at the middle of the light period under both photoperiods, with a broader and delayed profile under LD.

**Figure 10.**
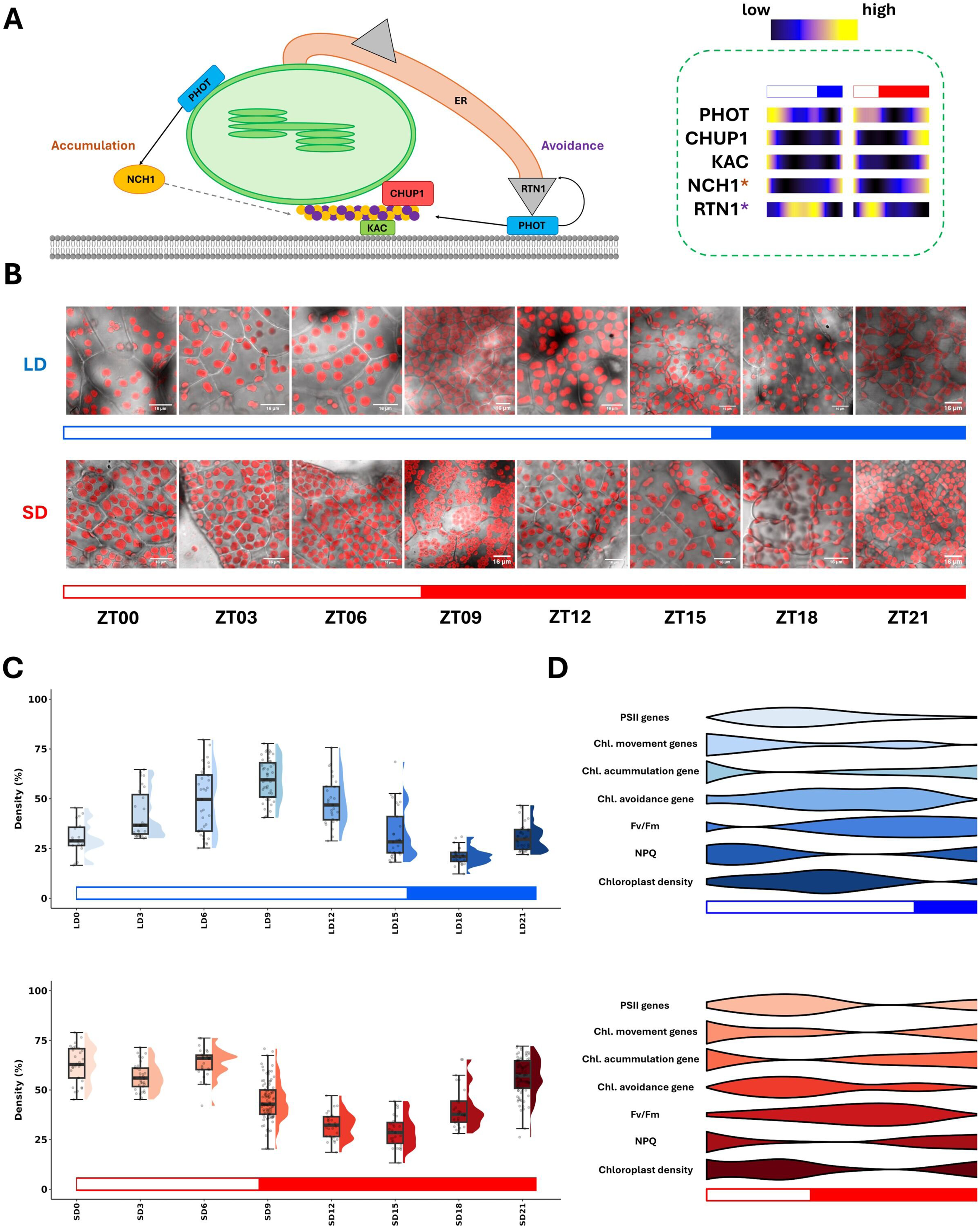
Photoperiodic timing of chloroplast photorelocation timing. **A)** Schematic representation of known mechanisms mediating chloroplast photorelocation in *Marchantia polymorpha* with heatmaps showing transcript abundance for photorelocation-associated genes under LD (left, blue) and SD (right, red) conditions. Purple asterisk marks the gene only associated with avoidance response, while orange one highlights an exclusive involvement in accumulation response. **C)** Representative confocal images showing exposed chloroplast area at each time point under LD (top, blue) and SD (bottom, red) conditions. Scale bars represent 16 µm in every image. **D)** Boxplots showing density of exposed chloroplast area in the periclinal cell walls, as percentage of total cell area occupied by chloroplasts under LD (top, blue) and SD (bottom, red). **E)** Violin plots integrating, from top to bottom, average expression of PSII genes, average expression of chloroplast movement genes (*MpPHOT*, *MpCHUP1* and *MpKAC*), average expression of accumulation-exclusive gene (*MpNCH1*), average expression of avoidance-exclusive gene (*MpRTN1*, *MpCHUP1* and *MpKAC*), Fv/Fm values, NPQ values and chloroplast densities in the periclinal cell walls under LD (top, blue) and SD (bottom, red) conditions.

Confocal microscopy revealed distinct chloroplast surface density at different time points, with a diel pattern (**Fig. 10B**). In agreement with transcriptomics and photosynthetic rate results, a strong rhythmic behavior was detected under both photoperiods, showing an evident anticipation under SD. Under LD condition, its peak occurred at ZT9 and its trough at ZT18, whereas under SD its phase was identified at ZT0 and its trough, 15 hours later (**Fig. 10C**). This implied a 3-hour anticipation in the rising patterns of chloroplast density under SD with respect to LD condition. The difference in peak width allowed presenting high chloroplast densities for light harvesting already at sunrise under SD and retaining them for a short time after sunset, to take advantage of the full photoperiod. Additionally, the density of chloroplasts exposed at daytime points was found to be significantly higher under both LD (p-value 1.1·10^-11^) and SD (p-value < 2.2·10^-16^) conditions when compared to the dark period. As for ZT0, under LD condition, chloroplast density values were significantly lower than those during the light period (p-value < 2.2·10^-16^) but no changes were detected with respect to those in dark (p-value 0.9523). In contrast, under SD condition, chloroplast density values were significantly higher at ZT0 than during the dark period (p-value < 2.2·10^-16^) and not lower than those during the light period (p-value 0.9987). Therefore, ZT0 appeared to be a transition point under LD condition, whereas under SD the transition took place earlier, during the dark period.

Finally, violin plots were used to compare the temporal profiles of photosynthetic physiology with their corresponding transcript expression to elucidate the coordination of the involved processes (**Fig. 10D**). Regarding chloroplast density, we observed individual patterns defined by the coordinated expression of the mentioned elements. The expression of *MpKAC* and *MpCHUP1* around dawn and late night under SD, combined with the expression of *MpPHOT* and the accumulation-related *MpNCH1*, both anticipating dawn, correlated with high chloroplast densities exposed in the periclinal cell wall from ZT0 onwards. Expression of *MpRTN1* peaked in the middle hours of the light period under these conditions (after ZT4), favoring the migration of chloroplasts to the anticlinal walls shortly thereafter. In the case of LD, the delayed expression of phototropin, motility proteins, and *MpNCH1* contributed to a delayed increase in periclinal chloroplast density. In addition, *MpRTN1* showed a more sustained peak over time and was delayed compared to SD, which could explain why, under long photoperiod, chloroplast density decreased later and less abruptly than under shorter ones. In terms of physiology, the results pointed to greater exposure of chloroplasts at the beginning of the day under SD, combined with a NPQ peak and a reduction in the PSII photochemical capacity to prevent photodamage during the transition to light, while genes encoding components of this complex peaked during the illumination period under both photoperiods.

## Discussion

In plants, diel rhythms control a wide range of metabolic and physiological processes. These rhythms exhibit significant seasonal variations, and parameters such as photoperiod are essential for their regulation. However, recent studies have reported notable differences in this regulation between vascular and non-vascular plants (Linde et al. 2017; Ferrari et al. 2019; Petersen et al. 2022; Tan et al. 2024). In this study, we have taken advantage of a design based on different photoperiods and free-running conditions to elucidate the seasonal plasticity of the dial and circadian transcriptome of Marchantia and its relationship with its photosynthetic physiology.

We detected a significant effect of photoperiod on the rhythmicity of the Marchantia transcriptome. 56% of the expressed transcriptome showed daily oscillations under LD condition, compared to 72% under SD. Arrhythmic genes showed very low expression and were associated with stress responses or cell cycle. Stress response genes may be expressed and show rhythmicity under specific induction conditions (Avelino et al. 2025). For their part, the low expression of cell cycle-related genes may be motivated by negative regulation of cell cycle under stress or by their localized expression in meristems, underrepresented tissues in the plant as a whole. Our results support those reported in other studies that indicate a lower percentage of rhythmic genes in bryophytes compared to chlorophyte microalgae (> 80 %) (Monnier et al. 2010; Zones et al. 2015; Romero-Losada et al. 2025) and higher than angiosperms such as *A. thaliana* (30% to 50%) (Bläsing et al. 2005), *Solanum tuberosum* (18 to 45%) (Hoopes et al. 2022), or *Oryza sativa* (41%) (Filichkin et al. 2011). However, these percentages decreased sharply under free-running conditions: only 25-30% of rhythmic genes cycled under LL and hardly ∼12 % did under DD. This is similar to the 15% expressed transcriptome reported under LL after ND entrainment (Lagercrantz et al. 2021), exactly 25-30% considering our estimates of rhythmic genes. LL-rhythmic genes showed involvement in photosynthetic and chlorophyll synthesis processes, while no major processes were enriched in DD-rhythmic genes. Thus, there existed a different regulation accounting for rhythm maintenance under LL and DD, and a predominant control of the circadian clock over photosynthesis in the presence of light that disappeared under constant darkness. Physiologically, this effect translated into higher RAE values in photosynthetic rhythms under DD than under LL observed in (Cuitun-Coronado et al. 2022). DD had a greater detrimental effect on the rhythmic transcriptome than LL under both photoperiods, similar to *A. thaliana* (Anwer et al. 2014), whose starch reserves were consumed during the night and did not allow rhythmicity to be sustained trough several days of DD (Graf et al. 2010). Genes oscillating under all conditions appeared to be involved in membrane integrity and chloroplast organization. Rhythmic profiles under free-running conditions showed significant differences from those under diel cycles. Both LL and DD reduced wave amplitude, consistent with desynchronization of autonomous circadian clocks of individual cells, already identified in other plants (Yakir et al. 2011). The observation of advanced and delayed phases under DD and LL, respectively, suggested a nocturnal organism (Hundahl et al. 2012; Imai et al. 2020). Most of gene expression was concentrated during the night, as advocated by the theory of light escape (Pittendrigh 1993). Radiation damage to DNA is magnified during the opening required for transcription, which is why many organisms perform this process during the dark period. These results also suggested that the dawn burst recently described in *A. thaliana* (Balcerowicz et al. 2021) was not present in this organism.

Diel gene phases organization showed significant differences between photoperiods, with SD exhibiting accumulation at two different time points, as observed under ND (Tan et al. 2024). However, LD showed only one significant accumulation at dusk. It is well known that diel rhythms are the result of combining light/dark response patterns with the anticipatory output of the circadian clock (Mehta et al. 2021). Breaking down these components, we have found that these whole transcriptome profiles correspond to a combination of response waves and a circadian wave. Circadian wave anticipated sunrise with one accumulation between ZT16-ZT20 under both photoperiods and another shortly after sunrise under LD. On the other hand, we observed rapid activation at dusk of dark-responsive genes, while light-responsive genes showed slow activation after dawn. This led to an overlap of the phase accumulations under LD, while in SD they became separated, giving rise to the observed bimodal profile. In addition, response to light or darkness did not depend on the photoperiod, since the identity of genes activated or repressed by these factors remained the same under both conditions.

Rhythmic genes presented greater amplitude and lower mesor under long photoperiod, albeit to a limited magnitude. On the other hand, a notable anticipation of phases was detected under SD condition compared to LD. This effect has been associated with a photoperiod-responsive gene expression wave and a fixed wave in marine phytoplankton, observable as two distinct waves under LL and DD conditions (Romero-Losada et al. 2025). This scenario was not observed in Marchantia, pointing to a more complex regulation, consistent with the greater number of circadian clock components in this species (Linde et al. 2017). In turn, SD condition also promoted the appearance of daily double peaks in gene expression, as has been described in species such as *O. tauri* (Romero-Losada et al. 2025). Biological processes and metabolic pathways showed advanced expression of the corresponding components under SD condition. Most of them, however, were clustered towards the nocturnal accumulations of phases described at the global transcriptome level. This could be due to a preference for transcription in darkness to avoid DNA photodamage. It is then expected that different biological processes would exhibit different delays in translation and post-transcriptional regulation to perform each one in its corresponding time window. However, there existed a group of biological processes whose expression did not follow these global dynamics. This was the case for photosynthetic processes expressed during the light period. In this instance, minor delays associated with these pathways and greater importance of transcriptional regulation over them would be expected. Other processes with cyclic behavior different from that of the mass of rhythmic transcripts were Calvin cycle, spliceosome and ribosome assembly, and protein processing. In this study, these temporal patterns have been linked to DNA-binding motifs to propose possible orchestrators of diel and circadian gene expression. The differentiated behavior of bZIP motifs such as the G-box and their rich presence in the promoters of morning-phased genes, LL-rhythmic genes, but not light-responsive genes, suggested a potential role for bZIP TFs in anticipatory diel regulation. In turn, these motifs were present in promoters of genes related to processes such as photosynthesis and light harvesting. Therefore, the circadian component appeared to exert a stronger influence on gene expression during dawn-morning, whereas light- and dark-responsive mechanisms governed gene expression at midday and during the night, respectively.

Core circadian clock components in Marchantia are a subject of debate. Due to its lack of a *CCA1/LHY* ortholog, the presence of orthologs of other proteins of *A. thaliana* clock has been studied (Linde et al. 2017). Although many of those proteins exist in the liverwort, studies on their expression have pointed to weak rhythms under free-running conditions (Lagercrantz et al. 2020). This is true even for *MpTOC1*, *MpPRR*, and *MpRVE*, proposed as a conserved clock loop. Other non-targeted studies have pointed to the potential importance of proteins such as *MpDET1,* based on their rhythmic pattern under LL following ND entrainment (Lagercrantz et al. 2021). In our study, the use of different entrainment photoperiods combined with both LL and DD conditions has allowed us to finely filter possible candidates. It pointed to a potential involvement of two Trihelix TFs and, interestingly, *MpHY5,* as mediators setting the rhythm of the circadian transcriptome, given their oscillations under all conditions. This discovery adds to the significant enrichment of G-box motifs recognized by *MpHY5* in LL-rhythmic genes but not in light- or dark-responsive genes. It is described how in *A. thaliana DET1* inhibits *HY5* activity (Cañibano et al. 2021) and activates the expression of *PIF4* (Gangappa and Kumar 2017). In Marchantia, we found that high abundances of the *MpDET1* transcript correlated with minimal expression of *MpHY5*. The alternating maximums in *MpHY5* and *MpPIF* expressions, also rhythmic under LL, suggest an antagonism between them, similar to that described in *A. thaliana* (Gangappa and Kumar 2017). This could serve to control the diel expression of their targets by binding to G-box in the promoters of the genes they regulate. The expression profile of genes with this motif revealed activation during the expression window of *MpHY5* and inhibition during that of *MpPIF*. All this leads us to propose a model in which the role of *MpDET1* at the beginning of the day is to reduce the expression of *MpHY5* and to promote that of *MpPIF*, coinciding with the increased levels of the latter and the expression decrease for G-box-regulated genes shortly after. In this case, photoperiodic regulation of these components anticipates their expression under SD condition. Finally, *MpDET1* and EE-containing-genes expression patterns ruled out the possibility of *CCA1*/*LHY*-independent repression of these targets at the beginning of the day (Lagercrantz et al. 2021). Indeed, their expression profiles supported the activating role of *MpRVE*, that also recognizes this motif and was expressed at the last hours of the light period. These differences in clock architecture or output with respect to flowering plants coincide with observations that decouple regulations typically linked to the circadian clock from the components studied. Thus, it has been shown that EC does not interact with *MpPIF* to regulate growth (Lagercrantz et al. 2022), unlike what occurs in *A. thaliana* (Nusinow et al. 2011). Other processes, such as the measurement of daylength for growth-phase transition, also do not appear to be mediated by the clock in this liverwort (Kanesaka et al. 2023).

As mentioned above, photosynthetic pathways exhibited very characteristic expression patterns that were strongly influenced by the photoperiod. Measurements of the photosynthetic physiology were integrated with its diel transcriptome to elucidate the temporal regulation of this process and the effect of the photoperiod over it. Fv/Fm represents the maximum capacity of plants to drive photochemistry when all PSII reaction centers are open (Murchie and Lawson 2013) and its variations may be related to the LHC themselves reorganizing to facilitate changes in NPQ and PSII reaction centers photochemical yield (Litthauer et al. 2015; Guardini et al. 2025). A shift in the maximum photochemical capacity profile was observed between photoperiods, advanced by 4 hours under SD condition, which allows for high photochemical activity to be maintained throughout the light period. In addition, rhythmic diel behavior was identified for non-photochemical quenching under both photoperiods, showing their maximums at dawn. NPQ protects photosystems against damage derived from the absorption of excess light by dissipating energy as heat (Zuo 2025). NPQ therefore increases in stressful situations, controlling PSII performance to prevent photoinhibition or photoinactivation. The gradual rise and fall in this parameter observed in this study has been related to light ramps, as opposed to on/off systems, which give rise to sharp NPQ spikes during the transition and minimum values at the rest of the time points (Lazzarin et al. 2024), unlike our results. Thus, under both photoperiods, the darkness-to-light transition seemed to impose a stress boundary that stimulated the activation of NPQ processes to cope with it. Observed Fv/Fm and NPQ complementary waves are associated with the possible sinks of PSII activation energy. Apart from losses due to basal fluorescence and photodamage, energy can be directed to either photochemistry or NPQ by LHC arrangement. An increase in the latter decreases the fraction of energy available for photochemistry and, therefore, may be indirectly linked to a decrease in the maximum photochemical capacity of PSII. On the other hand, the photorelocation of chloroplasts arranged these organelles on the periclinal surface at the beginning of the day under SD and towards noon under LD, thereby maximizing the hours of light capture under short photoperiods. This behavior points to diel regulation of the process, distinct from the response to sustained periods of darkness or varying light intensities previously reported (Komatsu et al. 2014). Anticipation in rhythmic profiles of motility and signaling genes involved in this process under SD pointed to a transcriptional regulation in addition to phototropin phosphorylation state (Kato et al. 2021; Noguchi et al. 2025), thereby allowing for tight seasonal control of light harvesting. Overall, Marchantia orchestrates the expression of its genes under diel cycles with the aim of maximizing light capture during the day under short photoperiods and restricting PSII activity at dawn to prevent photodamage. Conversely, under long photoperiods, it slows down light capture during the transition to light by restricting chloroplast area in periclinal walls, allowing it to delay photoprotection mechanisms and increase photochemistry in the final light hours, balancing the energy acquisition with the sustained stress over the extended light period.

## Methods

### Biological material and plant growth

*Marchantia polymorpha* plants (accession Takaragaike) were grown on Gamborg’s B5 medium from gemmae isolated from cups of mature thalli. Temperature was maintained at 20 °C and two different photoperiods were used, consisting of 16 h of light and 8 h of darkness for LD condition and 8 h of light and 16 h of darkness for SD condition, using 30 min ramps when turning the illumination on and off, up to a maximum irradiance of 100 μE m^-2^·s^-1^. Plants were grown for 4 weeks under SD condition and 3 weeks under LD condition to ensure a similar developmental state. To avoid positional bias, plates were collected randomly and their position changed during the growth period.

### RNA sampling protocol, purification and sequencing

Sampling was performed over the course of three successive days every 4 h, removing agar from thalli and freezing the biological material using liquid nitrogen. Nine plants per time point were pooled and grounded to fine powder. For free-running conditions, plants were transferred to constant light (LL) or constant darkness (DD) for two additional days after one day of acclimation (Sup. Fig. S1). Irradiance under LL condition was identical to the maximum reached under diel conditions. For the extraction of total nucleic acids, 1 ml of Trizol was added to 300 mg of powdered sample and, after homogenization, 200 μl of chloroform was added and centrifuged at 4°C for 15 min at 13500 g. 600 μl of phenol:chloroform:isoamyl alcohol were added to the same volume of supernatant, homogenized and centrifuged at 4°C for 10 min at 13500 g. 500 μl of the upper aqueous phase were collected and mixed with 500 μl of pure isopropanol, incubated on ice for 15 min and centrifuged as before for 10 min, after which the pellet was resuspended in 750 μl of 3 M LiCl and subjected to another 10 min centrifugation in the same conditions. The pellet was washed with 500 μl of 85% (v/v) EtOH and cold-centrifuged for 10 min. After discarding the supernatant, the resulting pellet was dried and dissolved in 21 μl of diethyl pyrocarbonate-treated (DEPC) water. Nucleic acid concentration and 260/280 and 260/230 ratios were quantified by Nanodrop ND-1000. Further RNA purification was performed using the Isolate II RNA Plant Kit (Bioline) following manufacturer’s instructions. Finally, isolated RNA was processed on an Agilent 2100 Bioanalyzer (DNA High sensitivity chip) to determine its quality.

cDNA libraries were prepared from 250 ng of RNA using Illumina TruSeq Stranded mRNA protocol following manufacturer’s instructions. Sequencing was performed on an Illumina NextSeq500 sequencer. Approximately 15 million 75 nt long single end reads were generated for each sample. Quality of the provided reads was analyzed using BaseSpace Onsite v 3.22.91.158. FASTQ Toolkit v1.0.0 software was used for filtering, trimming and de-multiplexing of data.

### RNA-seq data analysis

Quality of the resulting fastq files was assessed using FASTQC (Babraham Bioinformatics - FastQC A Quality Control tool for High Throughput Sequence Data). STAR (Dobin et al. 2013) and Samtools (Danecek et al. 2021) were used for mapping reads to genome version MpTak1v7.1 (Tanizawa et al. 2025). Transcript assembly and gene expression estimation as fragments per kilobase of exon and millions of mapped reads (FPKM) were carried out with StringTie2 (Kovaka et al. 2019) and R package Ballgown (Frazee et al. 2015). Exploratory analysis, noise correction and data normalization were performed using R packages factoextra (Kassambara and Mundt 2026), FactoMineR (Lê et al. 2008) and sva (Leek et al. 2012). R code developed for this analysis is available from the GitHub repository https://github.com/fran-romero-campero/CircaMarchantia.

### Analysis of rhythmicity and statistical comparative analysis of gene expression

R package CircaCompare (Parsons et al. 2020) was used to parametrically assess rhythmicity in expression patterns of genes adjusted to 24h- or 12h-period cosinusoidal waves to discern patterns consisting in one or two peaks per day, respectively. Estimation of wave parameters mesor, amplitude and phase allowed statistical comparison of differences in rhythmic profile between groups, applying a BH-adjusted p-value threshold of 0.05. Dampening of wave amplitude due to cell desynchronization under free-running conditions can hinder rhythmicity assessment. Because of this, conservation of rhythmic behavior after subjection to these conditions was determined by applying CircaCompare to a day of light/dark cycle and two days of free-running conditions, exclusively for rhythmic genes under diel cycles.

### Functional and circular gene set cohesion analysis

Due to the lack of an annotation package for *Marchantia polymorpha* in the Bioconductor repository, we generated a home-made annotation package for further functional analysis. To this end, the available information for MpTak1v7.1 (https://marchantia.info/) was processed using AnnotationForge to generate an R-interpretable package containing information about Gene Ontology (GO) terms, KEGG orthology, PFAM domains, SMART domains, PANTHER and KOG for each gene. It is available at https://github.com/fran-romero-campero/CircaMarchantia.

To test if gene phases associated with a certain functional set (for example, a GO category) were uniformly distributed around the circle or, conversely, they clustered towards specific diel time points, three different statistical tests were used. Rayleigh test is known to be the most powerful against uniformity against unimodal distributions (Landler et al. 2018). Additionally, Hermans-Rasson (HR) and Rao tests were used to identify multimodal rhythmic scenarios, overcoming the limitations of Rayleigh test (Landler et al. 2018, 2019). These tests were implemented in R packages circular and CircMLE (Fitak and Johnsen 2017). The resulting p-values were adjusted for multiple testing using BH method, and significance was determined at a 0.05 threshold. To identify cyclic gene sets with a different behavior from that of the global distribution of total cyclic mRNAs, a multivariate analysis of variance (MANOVA) was performed on the sines and cosines of the angles (gene phases) to linearize the circular data, as described in (Landler et al. 2021). This allowed for testing whether a sample was derived from the same distribution as the overall gene expression for the whole condition. The same approach was used to distinguish coordinated phase changes for gene sets between conditions. For sets that significantly changed their diel clustering, circular median was used as a predictor of the clustering phase in each condition. All measures were corrected by BH and the significance threshold was set at 0.05.

For transcription factor-binding sites (TFBS) analysis, DNA motifs in the region up to 1.5 kb upstream of the TSS of the genes were examined. This length was selected based on the reports of Marchantia having longer promoters than those of *Arabidopsis thaliana* (Romani et al. 2024). On the one hand, circular cohesion analysis was performed as described and enrichment of specific motifs for each discrete diel time point or in specific condition comparisons was determined using HOMER software v5.1 (Heinz et al. 2010) and its Plants database. The number of co-occurring motifs in the reduced sets was chosen based on database redundancy, selecting the minimum needed to represent each motif of interest: 17 for G-box and 12 for EE.

### Chlorophyll fluorescence measurements

Maximum quantum yield of PSII, Fv/Fm, and non-photochemical quenching (NPQ) were measured using an IMAGING-PAM M-Series chlorophyll fluorescence imaging system (Walz) following the protocol from (Casatejada et al. 2023). Plants were dark adapted for 20 min and then subjected to an induction-recovery curve program using 635 nm red actinic light at 81 μE m^−2^ s^−1^ for 7 min. 600 ms saturating pulses at 10,000 μE m^-2^ s^-1^ were applied every 40 s and their recovery in darkness was recorded for 8 min. Fv/Fm and NPQ parameters were calculated for four plants per time point using the equations of (Kramer et al. 2004). Fluorimetry files were processed using ImagingWinGigE V2.56p software in MAXI mode and statistical analysis was conducted using R package RAIN (Thaben and Westermark 2014).

### Confocal microscopy

To perform a morphological study of chloroplasts, a spectral Laser Scanning Confocal Microscope (Olympus FLUOVIEW FV3000) was employed. Excitation was performed with 488 and 561 nm lasers. Emission signals were measured within the green channel (500 to 540 nm, gain 504) and red channel (650 to 750 nm, gain 432). Images were recorded every 3 hours under both photoperiods. Image processing was performed using ImageJ software (Schneider et al. 2012) and rhythmicity statistics were calculated using R package RAIN. Comparison between light and darkness time points were performed using either Student’s T or Wilcoxon-Mann-Whitney tests based on data normality, assessed using the Shapiro-Wilk test.

### Data and Code availability

RNA-seq data generated in this study are freely available from the Gene Expression Omnibus database under the accession number GSE163368. The data analysis code developed using the statistical programming language R is freely available from the following GitHub repository CircaMarchantia: https://github.com/fran-romero-campero/CircaMarchantia.

## Supplemental Figures and Tables

**Sup. Fig. 1.**
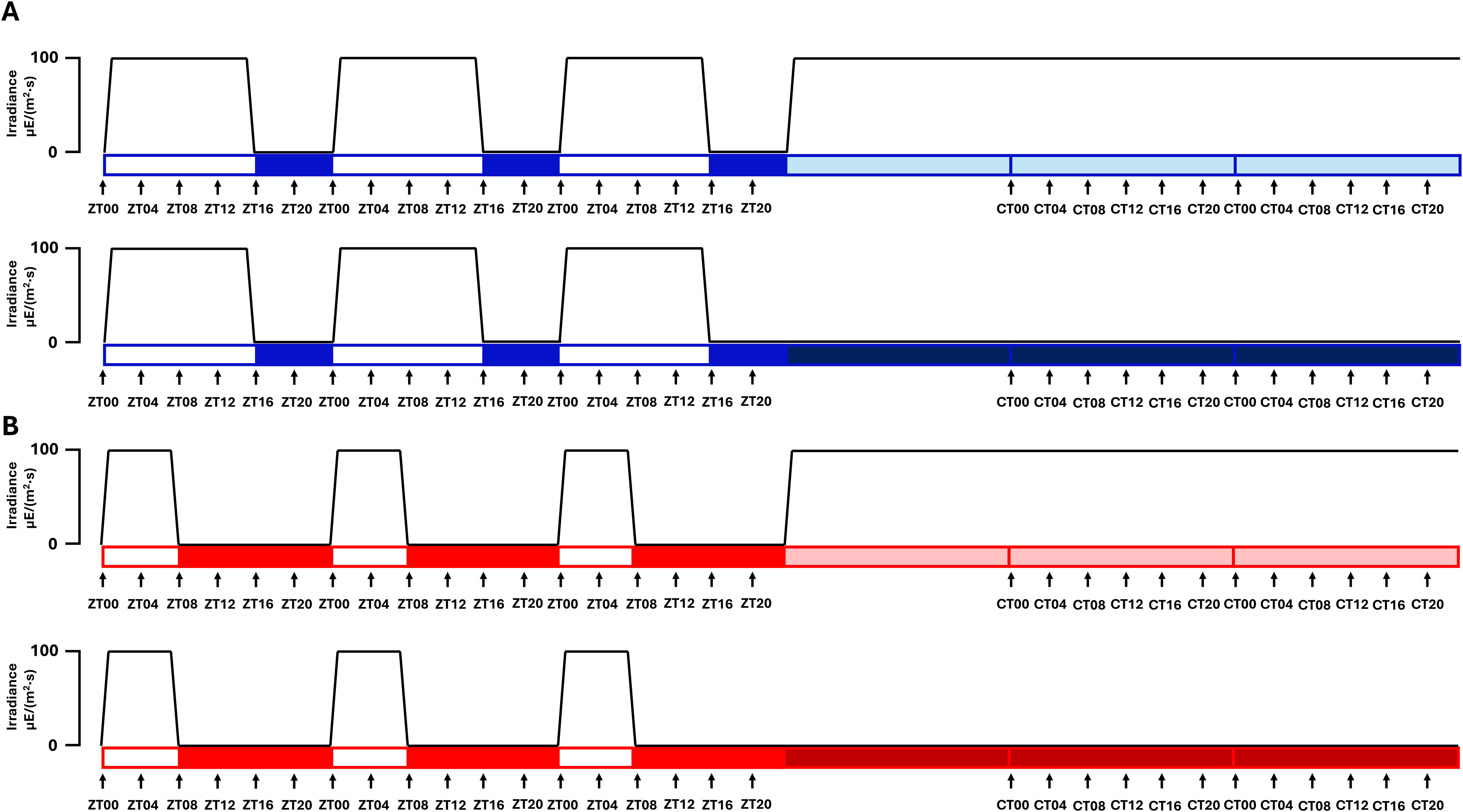
Experimental design. A) LL (top) and DD (bottom) after LD entrainment. B) LL (top) and DD (bottom) after SD entrainment.

**Sup. Fig. 2.**
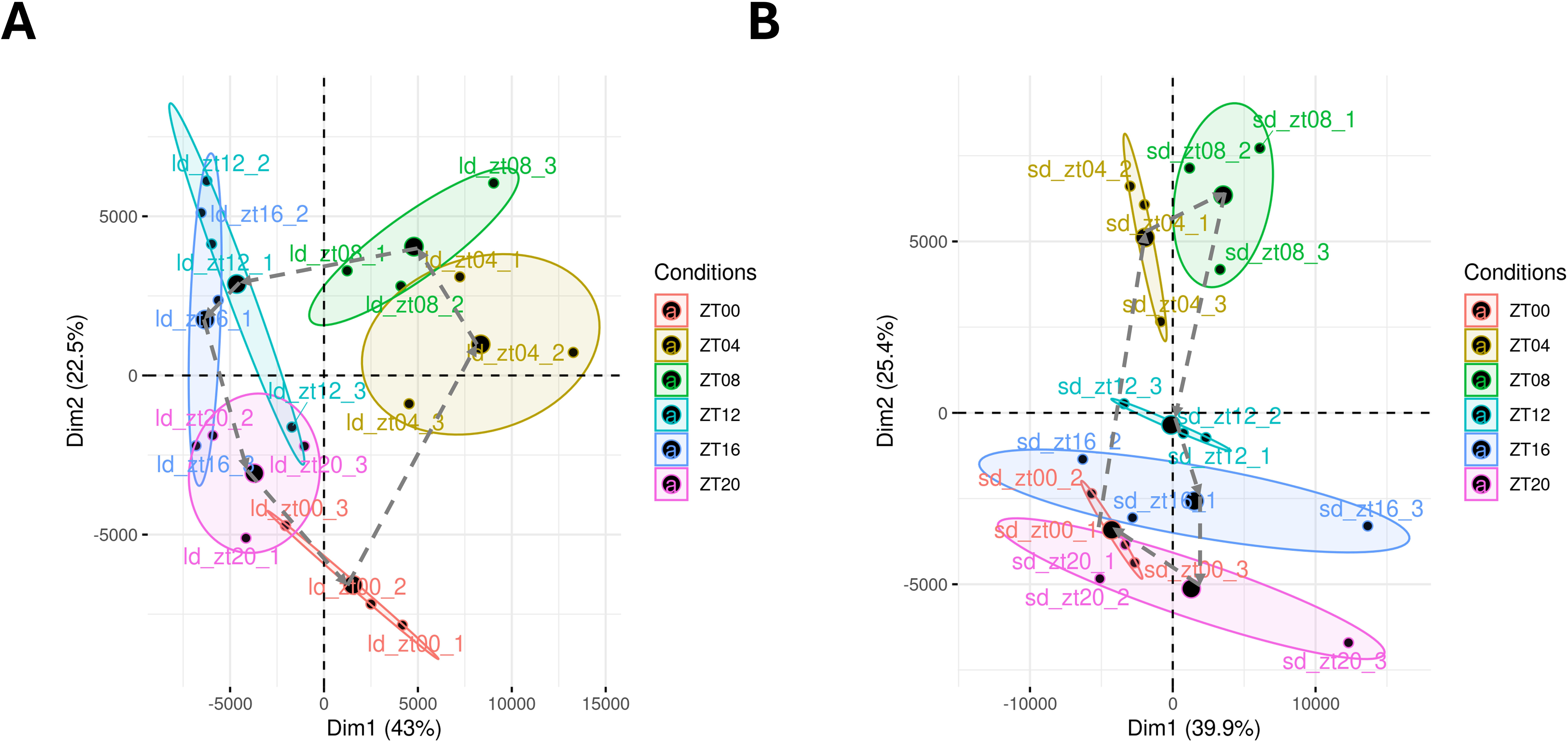
Principal Component Analysis of samples under **A)** LD and **B)** SD conditions. Small dots mark the projection of each sample in the 2D-space, while big ones correspond to the average for each condition. 95% confidence regions of each condition are represented by ellipses. Dashed arrows are used to indicate the temporal order of each condition.

**Sup. Fig. 3.**
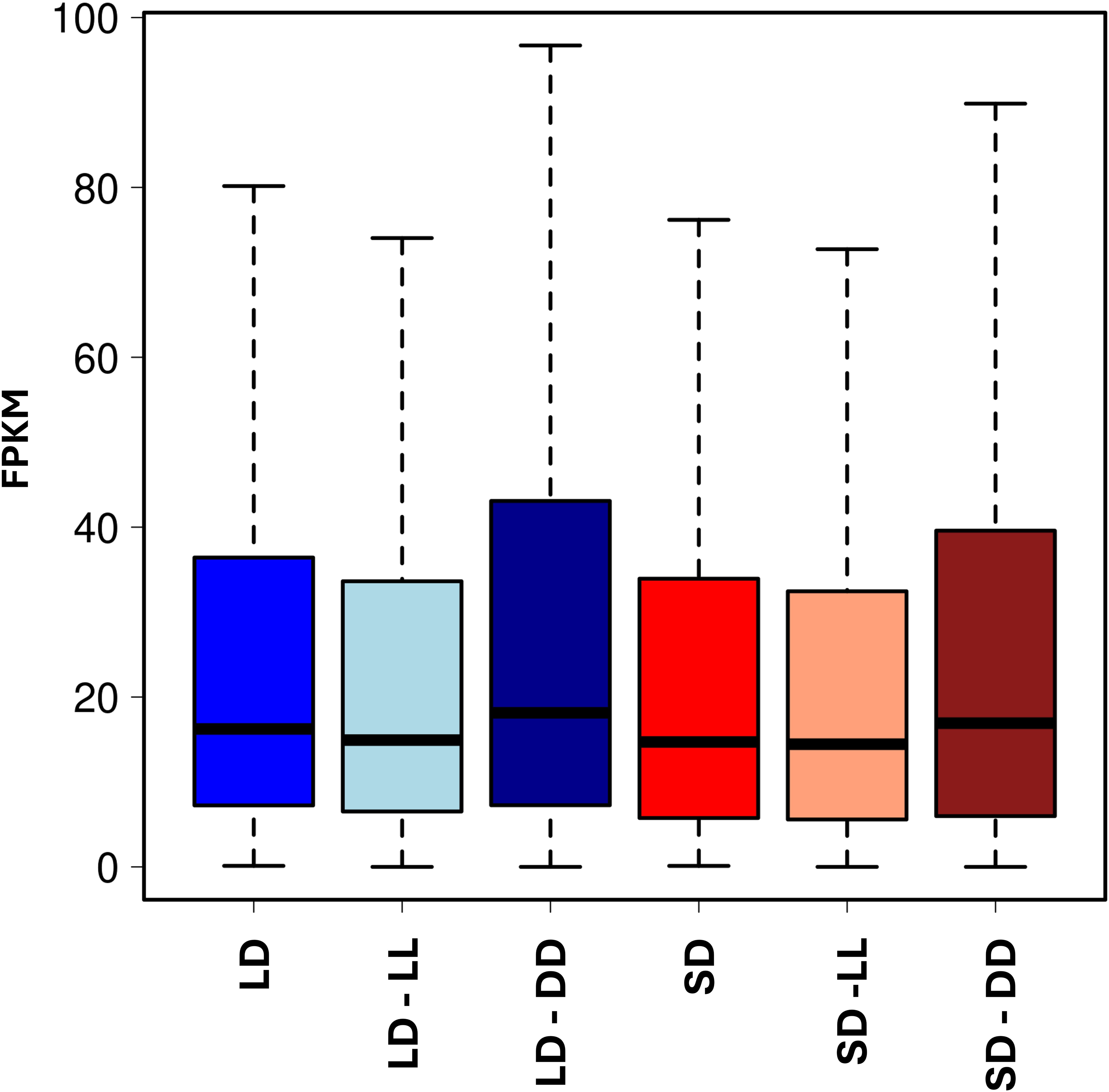
Boxplots showing gene expression by condition.

**Sup. Fig. 4.**
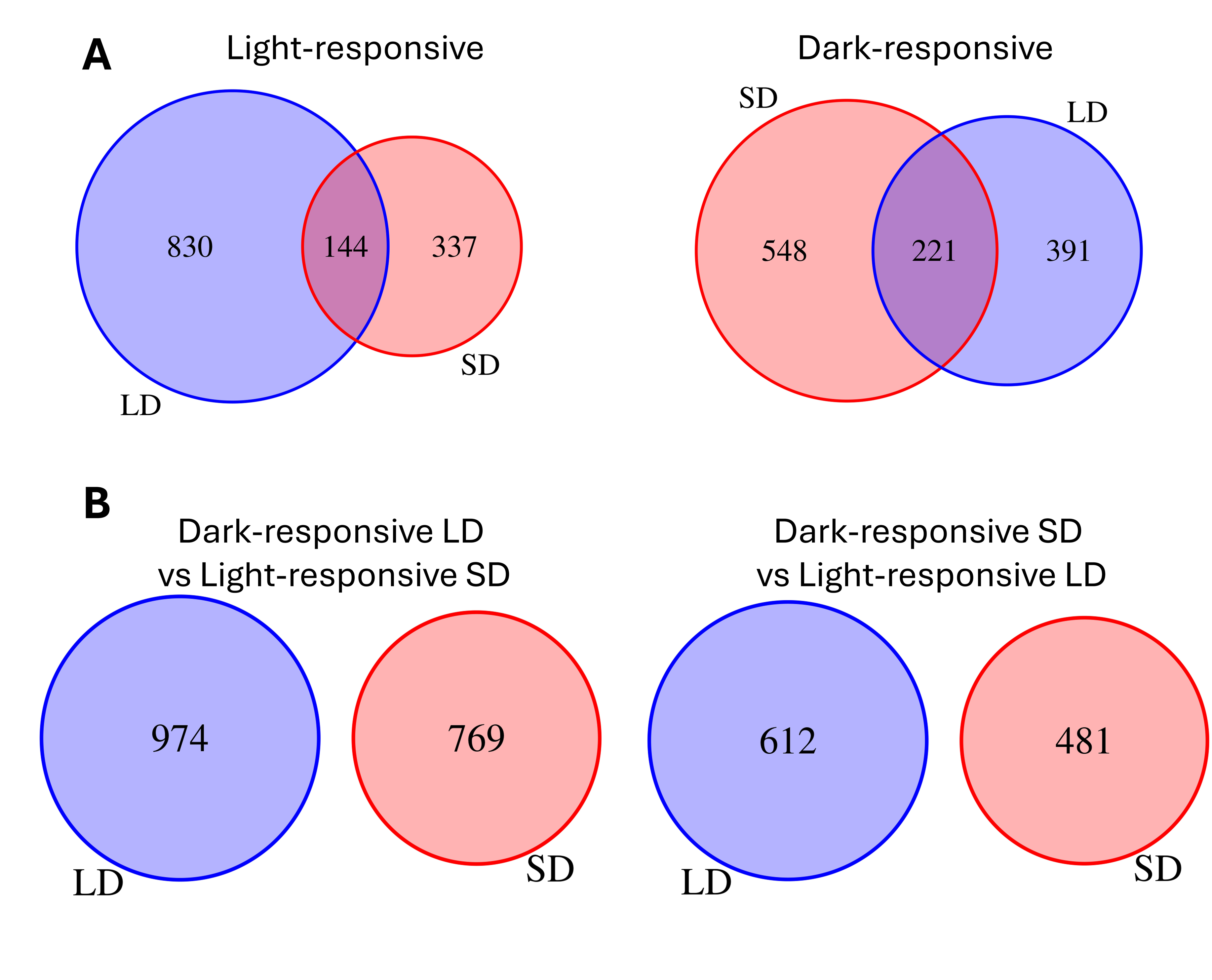
Venn diagrams of different rhythmic-genes sets. **A)** LD-light-responsive genes (blue) and SD-light-responsive genes (red). **B)** LD-dark-responsive genes (blue) and SD-dark-responsive genes (red). **C)** LD-dark-responsive genes (blue) and SD-light-responsive genes (red). **D)** LD-light-responsive genes (blue) and SD-dark-responsive genes (red).

**Sup. Fig. 5.**
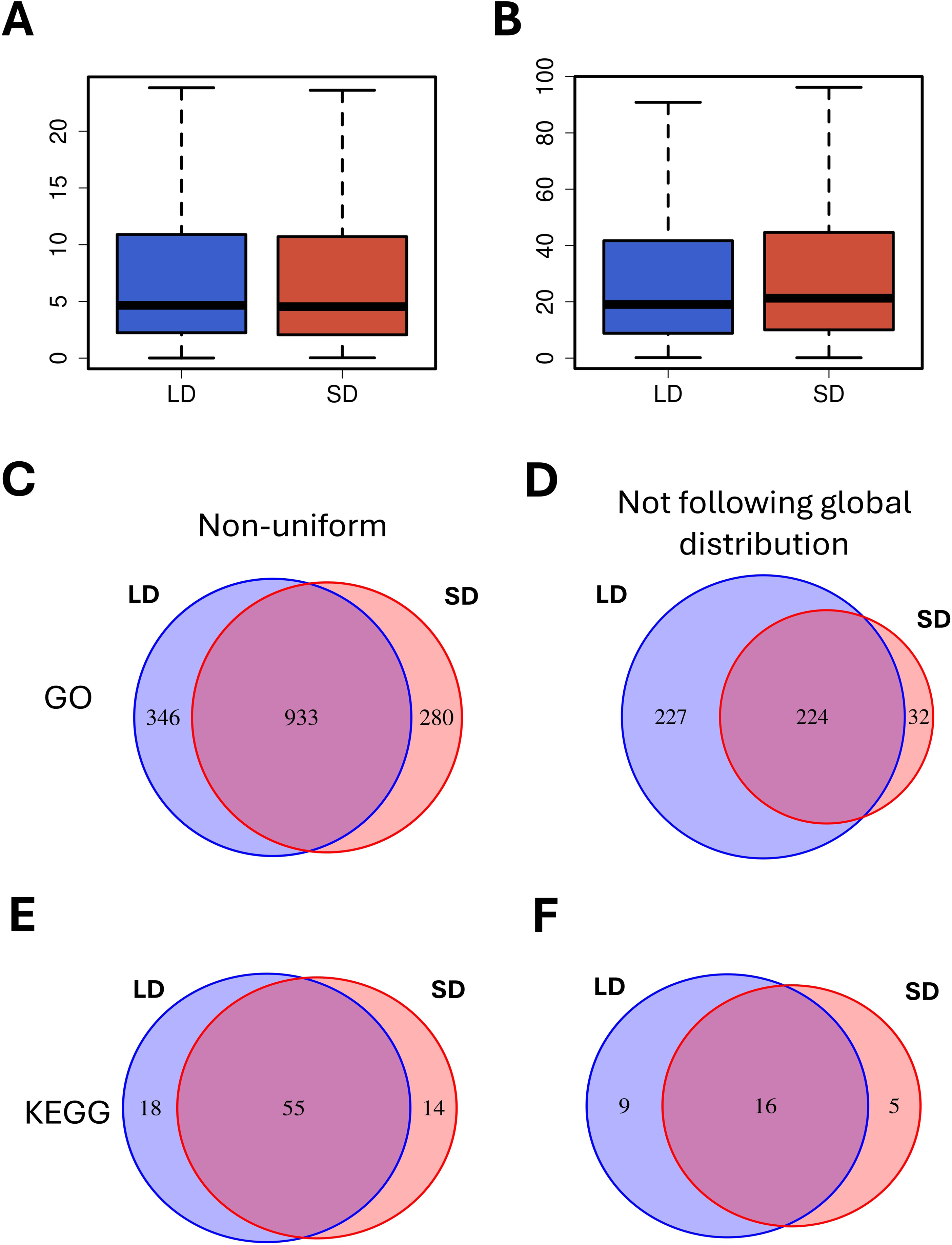
Venn diagrams of non-uniform and not experimentally distributed GO terms- and KEGG pathways-related gene expression by photoperiod. **A)** Boxplots representing amplitude of LD-(blue) and SD-rhythmic (red) genes. **B)** Boxplots representing mesor of LD-(blue) and SD-rhythmic (red) genes**. C)** Non-uniform GO terms under LD (blue) and SD (red) conditions. **D)** Non-experimentally-distributed GO terms under LD (blue) and SD (red) conditions. **E)** Non-uniform KEGG pathways under LD (blue) and SD (red) conditions. **F)** Non-experimentally-distributed KEGG pathways under LD (blue) and SD (red) conditions.

**Sup. Fig. 6.**
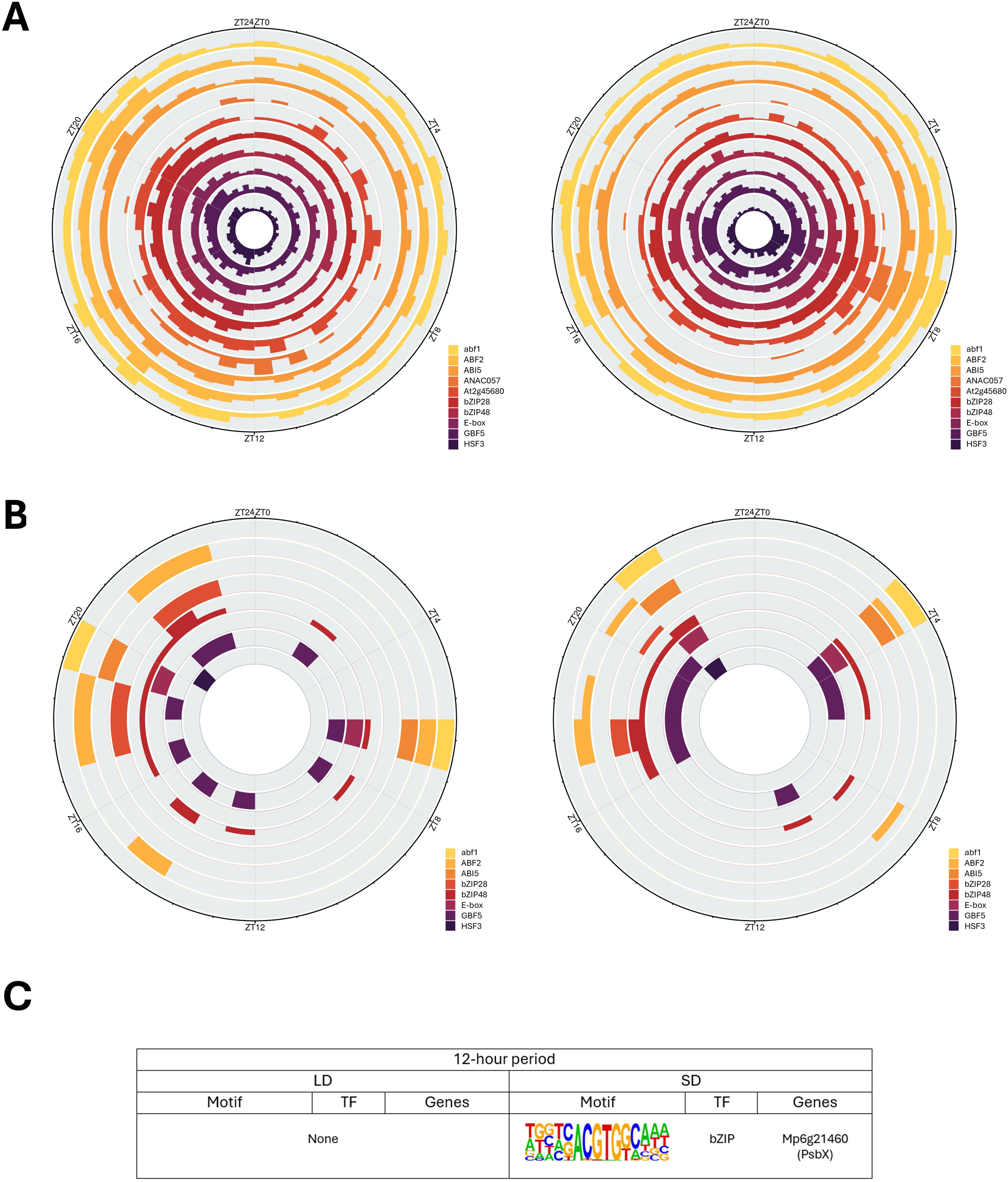
TFBS circular cohesion analysis and enrichment. **A)** Circular histogram of non-experimentally-distributed-TFBS-associated gene phases under LD (left) and SD (right) conditions. **B)** Circular histogram of non-experimentally-distributed-TFBS-associated gene phases under LD (left) and SD (right) conditions for genes showing a reduced motif set in their promoters (less than 20 motifs). **C)** TFBS enrichment for 12h-period-rhythmic genes when compared with the complete rhythmic-genes set under LD (left) or SD (right) conditions.

**Sup. Fig. 7.**
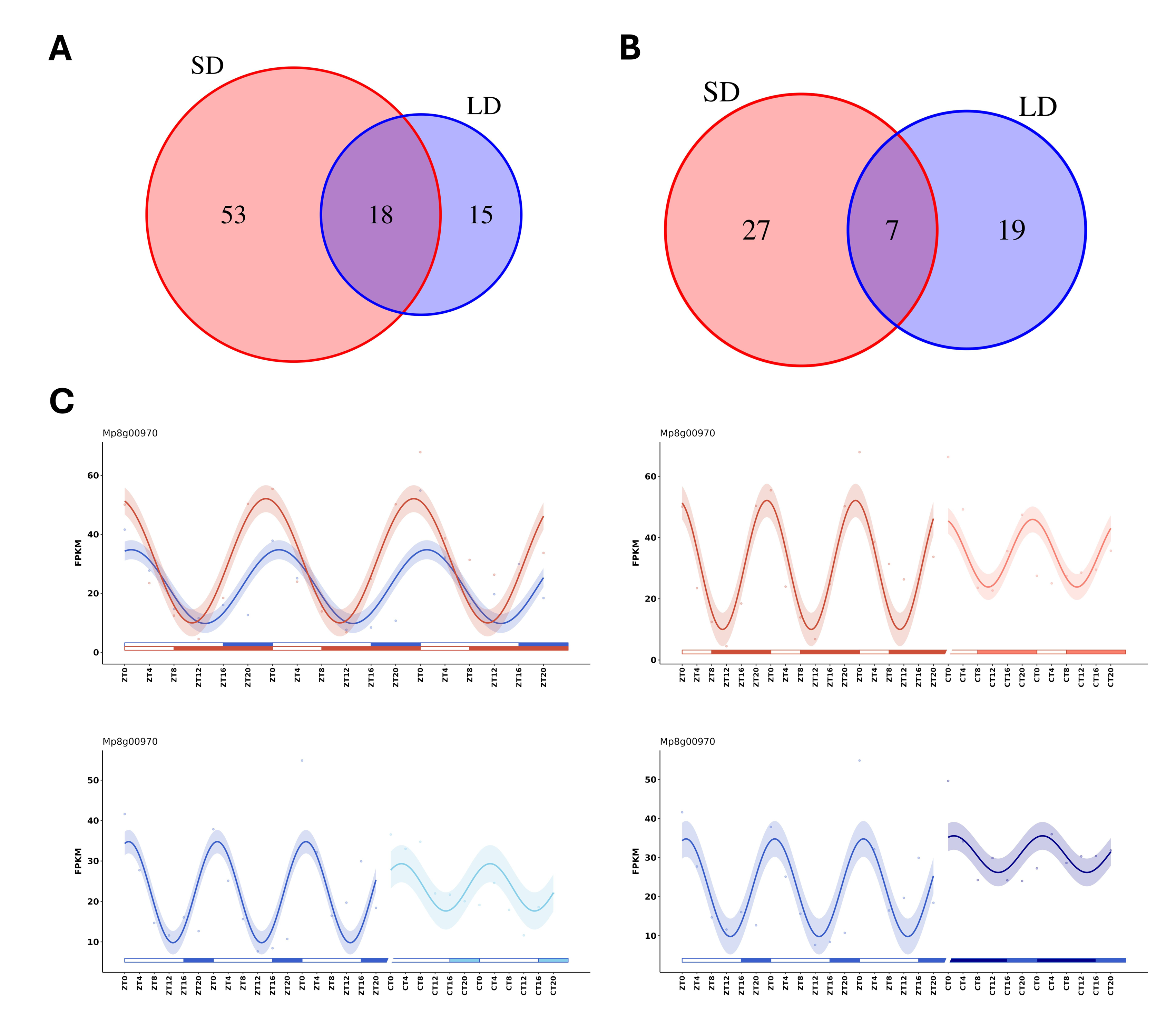
Venn diagrams and gene expression profile of rhythmic TFs. **A)** Venn diagram comparing LL-rhythmic TFs after entrainment in LD (blue) and SD (red) photoperiods. **B)** Venn diagram comparing DD-rhythmic TFs after entrainment in LD (blue) and SD (red) photoperiods. **C)** Gene expression profile of *MpDET1* in LD, LD-LL, LD-DD, SD and SD-LL.

**Sup Table S1.** GO terms enrichment for LD-LL-specific genes.

**Sup. Table S2.** GO terms enrichment for SD-LL-specific genes.

**Sup. Table S3.** Non-uniform GO terms in LD.

**Sup. Table S4.** Non-uniform GO terms in SD.

**Sup. Table S5.** Rhythmic GO terms not following the experimental distribution in LD.

**Sup. Table S6.** Rhythmic GO terms not following the experimental distribution in SD.

**Sup. Table S7.** Significantly phased-shifted BP GO terms with medians under LD and SD.

**Sup. Table S8.** Significantly phased-shifted MF GO terms with medians under LD and SD.

**Sup. Table S9**. Significantly phased-shifted CC GO terms with medians under LD and SD.

**Sup. Table S10.** Non-uniform KEGG pathways in LD.

**Sup. Table S11.** Non-uniform KEGG pathways in SD.

**Sup. Table S12.** Rhythmic KEGG pathways not following the experimental distribution in LD.

**Sup. Table S13.** Rhythmic KEGG pathways not following the experimental distribution in SD.

**Sup. Table S14.** Significantly phase-shifted KEGG pathways with medians in LD and SD.

**Sup. Table S15.** Rhythmic TFBS not following the experimental distribution under LD.

**Sup. Table S16.** Rhythmic TFBS not following the experimental distribution under SD.

## Acknowledgements

This work was supported by grants PID2021-123984OB-I00 (ELECTRA) and PID2024-158798OB-I00 (PERSEPHONE) from the Ministerio de Ciencia, Innovación y Universidades (MICIU)/Agencia Estatal de Investigación (AEI). M.R.G. was supported by an FPU predoctoral grant FPU22/00511 from the Ministerio de Ciencia, Innovación y Universidades.

## Author Contributions

M.R-G: formal analysis, investigation, methodology, validation, visualization, writing—original draft, review & editing. M.E.G-G: investigation, methodology, review & editing. J.D.B: investigation, methodology, review & editing. F.J.R-C.: conceptualization, formal analysis, investigation, validation, visualization, writing—original draft, review & editing. M.G-G: conceptualization, formal analysis, investigation, validation, visualization, writing—original draft, review & editing.

